# Seeing at Will: Shared Neural Representations of Motion Perception and Intention

**DOI:** 10.64898/2026.08.17.745279

**Authors:** Wentao Si, Eunhye Choe, Nathan H. Heller, Peter J. Kohler, Patrick Cavanagh, Viola S. Störmer, Peter U. Tse

**Author notes:** Please address correspondence to: Wentao Si.

## Abstract

Intention can bias perception in cases where two or more interpretations of a stimulus are possible. The neural mechanisms whereby intention influences perception are poorly understood. Here we investigated whether the intention to perceive horizontal or vertical motion in a bistable apparent motion stimulus pre-activates motion axes representations in sensory cortex before the position shift occurs and before motion in consciously perceived. Twelve participants underwent fMRI scanning under three conditions: (1) passively viewing quartet stimuli moving continuously along veridical trajectories; (2) passively viewing bistable apparent motion quartet stimuli induced by two pairs of squares alternately flashing at diagonally opposite corners of an invisible rectangle; or (3) while actively intending to perceive a single-shot apparent-motion stimulus as moving either vertically or horizontally. Multivariate decoding analyses revealed that activity patterns during the intention period of (3) generalized to patterns evoked by both (1) physical and (2) ambiguous motion perception. Cross-decoding was strongest within dorsal/lateral visual regions, including hMT+, V3AB, and the intraparietal sulcus, but was largely absent from ventral visual cortex. Widespread overlap was also observed between intention-related and perceptual motion representations throughout the dorsal/lateral visual cortex. Our findings suggest that intention may bias prospective sensory representations prior to the stimulus-driven processing that results in perceptual experience and that these representations closely resemble those associated with illusory motion perception. The fact that intention-related representations were mostly found within dorsal/lateral visual regions is consistent with top-down influences from attentional control systems.

**Significance Statement:** How thoughts and intentions influence perception is an important question in the cognitive neuroscience of consciousness. Using functional MRI and multivariate decoding techniques, we demonstrate that intention to see a subsequent instantaneous position shift as either horizontal or vertical motion recruits sensory representations that resemble those evoked during passive motion perception, with the strongest effects occurring in dorsal occipital and parietal cortex. These findings reveal how top-down processing proactively shape sensory representations and bias the perceived direction of subsequent motion, providing new insight into the neural mechanisms through which intention influences conscious visual experience.

## 1. Introduction

Humans are not merely passive recipients of their perceptual experience. Rather, conscious perception can be shaped by endogenous cognitive states such as expectation, attention, and intention. Attention selectively prioritizes relevant information and enhances behavioral performance for attended relative to unattended stimuli (Carrasco, 2011; Desimone and Duncan, 1995) and modulates neural responses across visual cortex when a target stimulus is presented (Pooresmaeili et al., 2010; Reynolds and Heeger, 2009; Treue and Martinez-Trujillo, 1999), and even when the stimulus is absent (Kastner et al., 1999; Serences & Kastner, 2014). Expectation can shape perception through predictions about likely upcoming inputs based on statistical regularities, sometimes allowing the brain to configure sensory processing even before a stimulus is present (Rao & Ballard, 1999; Kok et al., 2012; de Lange et al., 2018). Intention can also shape perception, but is distinguished by a goal-directed commitment toward a particular perceptual outcome, potentially biasing the weighting of subsequent sensory signals in advance. Deciding to see a bistable figure — such as the Necker cube — as facing one way rather than another illustrates how an active intention can determine which of two equally valid interpretations becomes conscious experience.

Many previous investigations of intention have focused on non-perceptual domains, demonstrating that neural activity preceding voluntary actions can shape the conscious experience of intending to act. The preparatory “readiness potential” precedes voluntary motor actions (Libet et al., 1983) and has been proposed to reflect the spontaneous accumulation of autocorrelated noise (Schurger et al., 2012). Similarly, intentions to perform mathematical operations can be decoded from distributed activity in prefrontal and parietal cortex before those operations are carried out (Haynes et al., 2007; Soon et al., 2013). These examples indicate that spontaneous neural processing can establish prospective neural states that bias subsequent processing. Comparatively less is known about how intention interacts with sensory cortices, and how intention, in pursuit of a goal, influences subsequent perceptual experience.

The apparent motion quartet offers a unique opportunity to study effects of intention on visual perception. The motion quartet is inherently bistable (perceived either as a pair of squares moving vertically or horizontally), while the stimulus itself is fixed: two dot pairs alternate in succession, with one appearing in the upper-left and lower-right positions, and the other in the opposite corners of an invisible rectangle (Ramachandran and Anstis, 1985). Psychophysical work has shown that intention — deliberately *choosing* to perceive vertical or horizontal motion — can bias the perceptual bi-stability of this ambiguous stimulus (Kohler et al., 2008; Sun et al., 2017). However, it remains unclear at what processing stage these effects arise, in particular whether top-down intention signals can modulate motion-direction-sensitive cortical regions such as hMT+.

Recent high-resolution fMRI work demonstrated that apparent motion evokes spatially dissociable column-like patterns within hMT+, comparable to those evoked by corresponding physical motion along the motion path (Schneider et al., 2019). Earlier findings showed that attended motion can be decoded from hMT+ (Kamitani and Tong, 2006), and that multivariate patterns of hMT+ BOLD signal activity predict perceived motion structure (Brouwer and van Ee, 2007). These findings raise the possibility that the intention to perceive a specific motion direction may generate neural activity patterns in motion-sensitive cortical populations even in the absence of a motion stimulus. Here, we isolated intention-related activity using a “one-shot” apparent-motion task, in which participants were cued to intend to see either horizontal or vertical motion during a motion-absent period preceding presentation of the ambiguous motion. We used multivariate decoding to identify encoding of intended perceptual outcomes, and cross-decoding to test if activity patterns associated with intention share a common representational format with those elicited during passive perception of physical motion and ambiguous apparent motion. Specifically, we asked whether frontoparietal control regions encode intended perceptual outcomes before one-shot apparent motion, whether motion-sensitive visual regions contain intention-related representations, and whether these preparatory neural states resemble the representations associated with subsequent conscious perceptual experience.

## 2. Methods

### 2.1 Participants

Eighteen participants (ages 18-62) from Dartmouth College and the surrounding community participated in the experiment. All had normal or corrected-to-normal vision and provided informed consent in accordance with protocols approved by the Institutional Review Boards at Dartmouth College. Due to the demanding nature of the study, which included a 1-hour pre-scan session and two to three scanning sessions totaling approximately 5 hours, several participants did not complete data collection. One participant was excluded from completing the experiment based on poor volitional control during pre-scan screening (see Text S1). Another three participants were not able to obtain reliable passive-viewing quartet aspect ratios using the pre-scan psychophysical procedures (see details under Stimuli and Procedure). Data from an additional participant were removed during the first scanning session because of poor fixation, and data collection for another participant was terminated after the participant fell asleep midway through the scan. The final sample consisted of twelve participants (9 males, 3 females), including authors N.H.H., E.C., and W.S. All participants except the authors were compensated with $10 for pre-scan psychophysics and $40 per hour for scanning sessions.

### 2.2 Stimuli, Apparatus and Procedure

All stimuli were created using PsychoPy v2024.2.4 (Peirce et al., 2019). The stimulus consisted of four white square inducers (1×1 dva) on a black background confined within a circular aperture of 3 degrees radius, and aspect ratio was varied by rotating inducer position around this aperture, since the perceptual threshold of the motion quartet depends on inducer eccentricity (Choe et al., 2025). To determine, for each participant, the individualized aspect ratio at which horizontal and vertical percepts were equally likely, we applied the method of limits with descending and ascending aspect ratio sequences to obtain a coarse estimate of the perceptual switch range, followed by the method of constant stimuli within this range to identify the point of subjective equality (Genç et al., 2011) (see Figure S7 demonstrating aspect ratio fit for all participants and Text S1for pre-scan procedures). Once the individualized aspect ratio was determined, this aspect ratio was used to present the quartet stimulus in Physical (i.e., continuous, analog motion) and Ambiguous (i.e., discrete) apparent motion and in the “one-shot” volitional control tasks. Prior to scanning, all participants except the authors (N.H., E.C., and W.S.) were screened for volitional control using a modified version of the volitional control task with the same number of trials as the in-scanner task. Participants with a hit rate >70% were eligible to proceed to scanning (see more details in supplementary text S1).

Participants completed two or three scanning sessions on different days. Physical and Ambiguous apparent motion passive viewing tasks were conducted in Session 1 or in Session 1 and Session 2, while the volitional control task was conducted in a separate session on a different day (Session 2 or Session 3) to minimize participants’ exposure to the volitional task while passive viewing. During scanning sessions, stimuli were projected onto a standing screen (18.9 dva by 11.8 dva projection area) outside of the MRI scanner bore (eye to mirror distance = 127mm; mirror to screen distance = 1160mm) using a Panasonic PT-DW750BU 7000-Lumen WXGA DLP projector. Eye movements were recorded using an Eyelink 1000+ (SR Research) that was placed inside the scanner underneath the screen (sampling at 1000 Hz). Physical and Ambiguous motion tasks were adopted from previous motion quartet studies (Schneider et al., 2019; Pizzuti et al., 2025) with minor modifications to reduce horizontal or vertical size biases regardless of individualized aspect ratios, as described above. A motion quartet stimulus was used to elicit illusory horizontal and vertical apparent motion percepts comparable to those elicited by the corresponding physical motion conditions with the same motion trajectories. The stimulus was identical to that used in pre-scan psychophysics. For the physical motion condition, the inducers translated at a constant speed either horizontally or vertically for 10-s blocks. Following four alternating blocks of horizontal and vertical motion (80-s total), a 16-s baseline condition was presented in which all four inducers flickered simultaneously without generating a motion percept. This 96-second block sequence was repeated six times per run. We added a 20-s fixation period at the beginning and end of each run, for a total runtime of 616 seconds (96 × 6 + 20 × 2). For the ambiguous motion condition, the overall timing and block structure were identical to the physical motion run, but the moving stimuli were replaced with a bistable apparent motion display. Specifically, pairs of diagonally opposite inducers were flashed for 150 ms (9 frames), followed by a 67 ms interstimulus interval (4 frames), producing a presentation frequency of 2.3 Hz that reliably induces apparent motion percepts (Schneider et al., 2019; Finlay and von Gru□nau, 1987). During scanning, participants continuously indicated whether they perceived horizontal (button 4, right thumb) or vertical (button 1, left thumb) motion using an MR-compatible button box. Each motion quartet run had a total duration of 10 min 16-s and consisted of 616 functional volumes acquired with a TR of 1 second. A total amount of 6 physical and 6 ambiguous motion runs were collected. To match physical and ambiguous motion tasks, participants were required to report vertical or horizontal perception in both types of runs.

The volitional control condition was performed on a different day. Each trial consisted of a pre-cue baseline period (16 - 20s), followed by an intention period (6 - 10s) during which participants voluntarily intended to perceive either horizontal or vertical motion as indicated by a brief 1s color cue (red or blue) in the beginning of the intending period. The color-to-intention mapping (e.g., red = prepare to perceive horizontal motion; blue = prepare to perceive vertical motion) was reversed halfway through the runs and conveyed to participants via instructions shown on the screen. Except for the cue, participants are presented with the same visual stimulation during pre-cue and intend period. This was followed by a switch inducing an instantaneous (one-shot) illusory apparent motion percept of either horizontal or vertical motion. Switched square pairs persisted for 4s after the switch and were followed by a response period (6s) in which participants reported their perceived motion direction. Response key mappings were randomly assigned to prevent motor preparatory contamination. Participants received a pseudorandomized mapping of horizontal and vertical percept on two of four buttons during the response period of every trial. Total trial duration was always 36 seconds and each volitional control run had 10 trials and a total duration of 6 minutes and consisted of 360 functional volumes acquired with a TR of 1 second. A total of 16 volitional control runs were collected during the last session. Across the 16 runs, half of the trials within each intention condition (horizontal or vertical) began with squares presented in the upper-left and lower-right positions, whereas the remaining half began with the opposite diagonal configuration. To avoid contamination from cue association, the color–intention mapping was reversed halfway through the experiment: blue indicated vertical intention and red indicated horizontal intention during runs 1–8, whereas red indicated vertical intention and blue indicated horizontal intention during runs 9–16. In addition, 10% of all trials were pseudo-randomly designated as catch trials to monitor for report validity, with equal numbers assigned to the horizontal and vertical conditions. During these trials, participants were forced to perceive a motion axis opposite to the instructed intention via the insertion of an intermediate stimulus square along the apparent motion trajectory. Each run contained 0, 1, or 2 catch trials. Please see Figure 1 for a schematic illustration of the experimental design.

**Figure 1.**
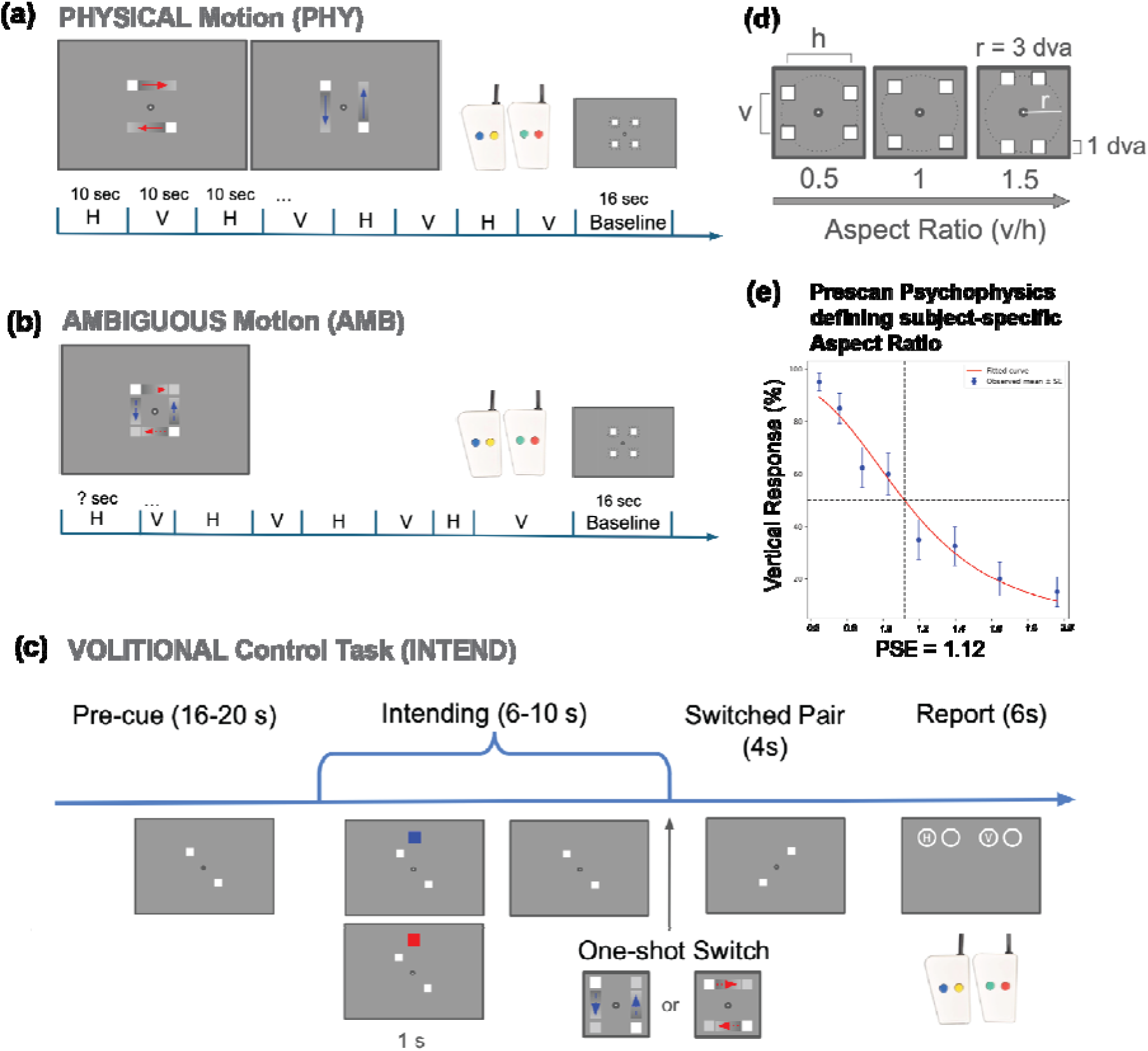
Experimental paradigm. (a–b) Experimental design for passive viewing tasks in the fMRI session. Participants viewed either physical (veridical) motion presented in alternating 10-s blocks of horizontal (H) and vertical (V) motion (a), or ambiguous motion stimuli where subjective percepts would vary continuously but at a random rate, followed by a baseline period where stimulus did not move. In both conditions, participants reported perceived motion direction via a button press. (c) Volitional control condition performed in a separate session. Each trial consisted of a pre-cue period, an intending period during which participants voluntarily intended to perceive a motion axis (horizontal or vertical) as instructed by a brief color cue, a switched-pair display inducing an instantaneous illusory motion axis, and a response period for reporting the perceived direction with sample key mapping across trials (d-e). Pre-scan psychophysics used to determine subject-specific stimulus parameters to reduce stimuli induced bias. A combination of the method of constant stimuli (to estimate ascending and descending aspect-ratio induced perceptual switch ranges) and the method of limits (to precisely determine the point of subjective equality) was used to derive each participant’s individualized aspect ratio prior to scanning. (d) The motion quartet was presented within a circular aperture of 3° radius, and each square subtended 1 × 1° of visual angle. (e) Example pre-scan result from one participant.

hMT+ localizer data were collected either in session 2 or session 3. The stimuli were also adopted from previous motion quartet studies (Schneider et al., 2019; Pizzuti et al., 2025) based on Huk et al. (2002). White dots (n = 200; dot size = 0.2° visual angle) were presented on a gray background and moved either inward or outward relative to the center of the aperture for 10-s at a speed of 8° visual angle/s. This motion period was followed by 10-s of stationary dots. The task–rest sequence was repeated 14 times. The total scan duration was 4 min 50 s (270 volumes; TR = 1 s). Observers were instructed to press a button immediately after detecting a subtle and brief change in the fixation color. The timing of 7 fixation color change events was pseudo-randomized within each run.

### 2.3 MRI acquisition and pre-processing

MRI data were acquired at the Dartmouth Brain Imaging Center using a 3 Tesla Siemens MEGNETOM Prisma scanner. At the beginning of each session, after a brief scout scan, pairs of three-dimensional *B*_0_ field maps with opposite phase-encoding directions (anterior-to-posterior and posterior-to-anterior) were collected to facilitate EPI distortion correction during preprocessing. Participants completed six physical-motion and six ambiguous-motion functional runs acquired using multiband T2*-weighted echo-planar imaging (EPI; TR = 1,000 ms; TE = 30 ms; 2.5 mm isotropic resolution; flip angle = 60°; multiband factor = 4; GRAPPA acceleration factor = 2). These runs were typically completed in a single scanning session but, when necessary, were continued in a second session. On a separate day, participants completed 16 functional runs of the volitional control task and three hMT+ localizer runs, acquired using the same EPI parameters. A high-resolution T1-weighted anatomical image was also acquired using an MPRAGE sequence (0.94 mm isotropic resolution; flip angle = 8°; TR = 2.3 s; TE = 2.3 ms).

Results included in this manuscript come from preprocessing performed using fMRIPrep 24.1.1 (Esteban et al., 2019, Esteban et al., 2018), which is based on Nipype 1.8.6 (K. Gorgolewski et al., 2011; K. J. Gorgolewski et al., 2018). fMRIPrep generates a description of the algorithms, software packages and procedures included in its pipeline. This description is reprinted in the supplementary materials (Text S2) with minor edits to clarify the field-map correction procedures in a 2-session layout.

The final three participants were scanned following a scanner software upgrade to Siemens XA60 that unexpectedly altered the default settings of two RF pulse parameters: the sinc excitation pulse bandwidth–time product (BWTP) was reduced from 5.2 to 1.0, and the fat-saturation flip angle was reduced from 110° to 30°, while other acquisition parameters were preserved. The reduced BWTP may broaden the effective slice profile and increase through-plane partial-volume effects and slice crosstalk, potentially degrading spatial resolution and adversely affecting multivariate decoding. We therefore repeated the searchlight analysis using only the nine participants scanned before the upgrade. The resulting searchlight decoding maps were highly similar to those obtained with the full sample (N = 12), with negligible differences in the spatial distribution of significant decoding clusters (Figure S9).

### 2.4 ROI definition

hMT+ was functionally defined using 3 hMT+ localizer runs as described above under stimuli and procedure. To analyze localizer data, we first fit a generalized linear model (GLM) to BOLD time series data from all runs at each vertex in *fsnative* space using *Nilearn* (v 0.12.0), with two regressors of interest estimating brain responses to stationary and moving dots (modeled with a canonical hemodynamic response function). Besides the regressors for each condition, we used nuisance regressors for motion (six rigid motion directions estimated during preprocessing) and for scanner drift (first- and second-order polynomials). The model was corrected for temporal auto-correlation (AR2). We took the contrast between moving and stationary dot periods and set a threshold corrected for multiple comparisons using false discovery rate; q(FDR) < 0.05. hMT+ for each subject was delineated manually using the Freesurfer GUI within the significant cluster around the ascending limb of the inferior temporal sulcus with the help of Wang atlas TO1 and TO2 contours plotted on individual *fsnative* surfaces as reference. Surface hMT+ ROIs for each subject were then projected back to their respective volumetric space using customized scripts, AFNI version 25.2.05 (Cox, 1996) 3dcalc and 3d*Surf2Vol*.

Beyond functionally defined hMT+, regions of interest (ROIs) in the dorsal/lateral and ventral visual hierarchy were defined using the probabilistic atlas of topographically organized visual areas created by Wang et al. (2015). The atlas was generated from functional localizer data collected in approximately 50 participants and includes 25 topographic ROIs spanning 22 visual areas. Individual cortical surfaces were standardized using icosahedral tessellation and surface-based projection methods described by Argall et al. (2006) after which the probability of each surface vertex belonging to a given ROI was estimated across participants. ROIs were defined using a maximum-probability approach, whereby a vertex was assigned to a visual area only if it was more frequently classified within that ROI than outside it across participants. In such cases, the vertex was labeled according to the most probable ROI identity. This probabilistic atlas preserves the large-scale organization of individually defined retinotopic regions and generalizes well to independent participants who were not included in atlas construction. The atlas was obtained from https://napl.scholar.princeton.edu/resources. ROIs were transformed from the standardized surface space into each participant’s native FreeSurfer surface space (*fsnative*) using the conversion scripts provided by Takemura and Benson at the same resource. We then converted surface ROIs into individual volumetric space registered to their T1w image using customized scripts, AFNI version 25.2.05 (Cox, 1996) 3dcalc and 3d*Surf2Vol* for further analyses. We excluded four ROIs from our analysis, IPS4, IPS5, SPL, and FEF, each of which covered a comparatively small number of vertices/voxels in the probabilistic map. To mitigate potential imprecision associated with atlas-based ROI definitions, several neighboring visual areas were combined into larger composite ROIs using custom scripts in our analysis.

Specifically, V1, V2, and V3 were combined to form early visual cortex (V1-3); V3A and V3B were combined as V3A-B; IPS0 and IPS1 were grouped as IPS0-1; PHC1 and PHC2 were combined as PHC1-2; IPS2 and IPS3 were grouped as IPS2-3; LO1 and LO2 were combined as LO1-2; and TO1 and TO2 were combined as TO1-2 (corresponding to Wang hMT+). For comparisons of decoding performance between dorsal and ventral visual cortex, decoding performance was first calculated separately for each individual, uncombined atlas-defined ROI and then averaged across the ROIs assigned to each division (dorsal or ventral) within each participant. The dorsal–ventral boundary was defined by the horizontal meridian in V1 defined by the Wang atlas. Accordingly, V1, V2, and V3 were subdivided into their dorsal and ventral components (V1d/V1v, V2d/V2v, and V3d/V3v, respectively) rather than treated as combined ROIs.

### 2.5 Univariate Analyses

Univariate analyses were conducted to identify cortical regions showing differential activity during the intention period relative to the pre-cue baseline period. Functional data were first preprocessed and normalized to the MNI152NLin2009cAsym standard space. A general linear model (GLM) was fit using concatenated runs from the intention task. Intention and pre-cue baseline periods were modeled as separate regressors convolved with a canonical hemodynamic response function (HRF), irrespective of intended motion direction (horizontal or vertical). Temporal autocorrelation was modeled using a first-order autoregressive (AR1) model. Six motion parameters, along with linear and quadratic scanner drift terms, were included as nuisance regressors. Subject-level contrast maps for the intention > pre-cue baseline comparison were entered into a second-level one-sample *t*-test implemented in *Nilearn* (0.12.0). Spatial smoothing using a 6-mm full-width at half-maximum (FWHM) Gaussian kernel was applied within the second-level model prior to group statistical inference. Group-level *t*-maps were subsequently projected to *fsaverage* surface space for visualization. Multiple-comparison correction was performed using false discovery rate (FDR) correction at *q* < 0.05 (Figure 3b; Figure S2).

To further characterize the temporal dynamics of activity during the intention task in retinotopic visual regions, z-scored BOLD time-courses were extracted from ROIs based on the probabilistic atlas of Wang et al. (2015). Time-courses were aligned either to cue onset or to the instantaneous motion switch. Because the delay period contained a 6–10 TR temporal jitter, only 6 TRs following cue onset and 6 TRs preceding the instantaneous switch were visualized (Figure 3a; Figure S3). Differences between horizontal and vertical intention trials at each TR were assessed using paired statistical comparisons across participants, with FDR correction applied across all time points across all ROIs (*q*FDR < 0.05).

### 2.6 Multivariate Analyses

Trial-wise samples used in ROI-based multivariate analyses were obtained by fitting a general linear model (GLM) separately for each run using *Nilearn* (version 0.12.0). For all three run types (physical motion perception, ambiguous motion perception, and intended motion perception), horizontal and vertical periods were modeled as separate regressors using a canonical HRF. Temporal autocorrelation was modeled using a first-order autoregressive (AR1) model. The GLM additionally included six motion parameters as well as linear and quadratic scanner drift terms as nuisance regressors. Single-trial beta values were then extracted for each voxel within each ROI in individual anatomical space and subsequently used as samples for ROI based multivariate decoding analyses. Additionally, the same GLM procedures were applied in *fsaverage* surface space (sub-sampled *fsaverage5* for each subject to reduce computational cost) to estimate single-trial beta values for each cortical vertex, which were subsequently used for whole-cortex searchlight analyses.

All decoding analyses were performed using linear support vector machines (scikit-learn Python package v1.7.1) classifying two classes (horizontal vs. vertical) with default regularization parameter (c = 1.0) in customized scripts. Model performance was evaluated with ROC-AUC score to control for slight class imbalance due to eliminating ‘miss’ intention trials (i.e. seeing motion axes orthogonal to the instructed intention). Within-domain decoding performance was estimated separately for each subject using a leave-one-run-out cross-validation procedure. For cross-domain decoding analyses, classifiers were trained on all but one run from one condition and evaluated on all runs from a different condition. This was done for each run. Decoding performance was computed separately for each subject, and group-level performance for each train–test pair was quantified as the mean decoding accuracy across participants.

To assess whether group-level decoding performance was significantly above chance (50%), training labels were pseudo-randomly shuffled 1,000 times for each subject to generate subject-level null distributions. A group-level null distribution was then constructed using Monte Carlo sampling: for each iteration, one value was randomly sampled with replacement from each subject’s null distribution, and the mean across participants was computed. Repeating this procedure 100,000 times produced a group-level null distribution centered at chance. Empirical p-values were computed by comparing the observed group mean decoding accuracy against the group-level null distribution generated through Monte Carlo sampling. Empirical p-values obtained from probabilistic atlas ROIs decoding analyses were corrected for multiple comparisons using the Benjamini–Hochberg false discovery rate (FDR) procedure across 81 comparisons across 9 ROIs and 9 conditions (3 within domain tests and 6 across domain tests).

When testing differences in representational strength between physical motion, ambiguous motion, and intention conditions, group-level null distributions for each ROI were computed by pairwise subtraction of the corresponding group-level null distribution between conditions (e.g., physical null – ambiguous null). Empirical *p*-values were then computed by comparing the observed group mean difference against the corresponding group-level difference null distribution. Multiple comparisons were corrected using the Benjamini–Hochberg false discovery rate (FDR) procedure across 27 comparisons, corresponding to 3 condition-difference contrasts across 9 ROIs. Additionally, to ensure fair comparisons across conditions (i.e., that stronger decoding performance in the physical motion condition was not driven by a larger number of samples), horizontal and vertical trials were subsampled and balanced across physical, ambiguous, and volitional runs according to the condition with the smallest number of trials for each subject. All within-domain decoding and performance comparisons are trained and tested within sub-sampled, balanced samples.

For comparisons of decoding performance between dorsal and ventral visual regions, decoding performance was first averaged across the individual, uncombined atlas-defined ROIs assigned to each cortical division (dorsal and ventral) within each participant. The corresponding permutation distributions were averaged across the same ROIs, yielding participant-level dorsal and ventral null distributions. Paired dorsal–ventral differences were then computed for both the observed decoding performance and the corresponding permutation distributions. To generate the group-level null distribution, one value was randomly sampled from each participant’s null difference distribution and averaged across participants; this procedure was repeated 100,000 times. Two-sided empirical *p* values were calculated as the proportion of group-level null differences whose absolute magnitude was equal to or greater than that of the observed group-mean difference. *P* values were corrected for multiple comparisons using the Benjamini–Hochberg false discovery rate (FDR) procedure across all train–test pairs, with significance assessed at *q*FDR < 0.05.

Although the original design balanced left- and right-tilted square pairs across horizontal and vertical intention conditions during the intention period of the volitional control task, missed trials (see Figure 2a) introduced unintended imbalances in the distribution of left- and right-tilted square pairs across the two classes. This raised the possibility that the classifier could decode low-level retinotopic signals associated with the tilted square pairs, rather than top-down intention-related signals, during within-domain volitional decoding. To mitigate this issue, for each cross-validation iteration, the numbers of left- and right-tilted square pair trials were balanced within the training set, while classifier performance was evaluated on the full set of trials in the held-out run. Within-domain decoding analyses using the full, unbalanced set of collected samples are reported in the Supplementary Materials (Figure S5). No qualitative differences were observed between within-domain decoding results obtained using balanced subsampled trials (Figure 5a) and those obtained using the full unbalanced samples (Figure S5). We therefore concluded that sample size differences between conditions in our data did not qualitatively impact decoding performance and opted to use full samples in subsequent searchlight decoding analyses.

**Figure 2.**
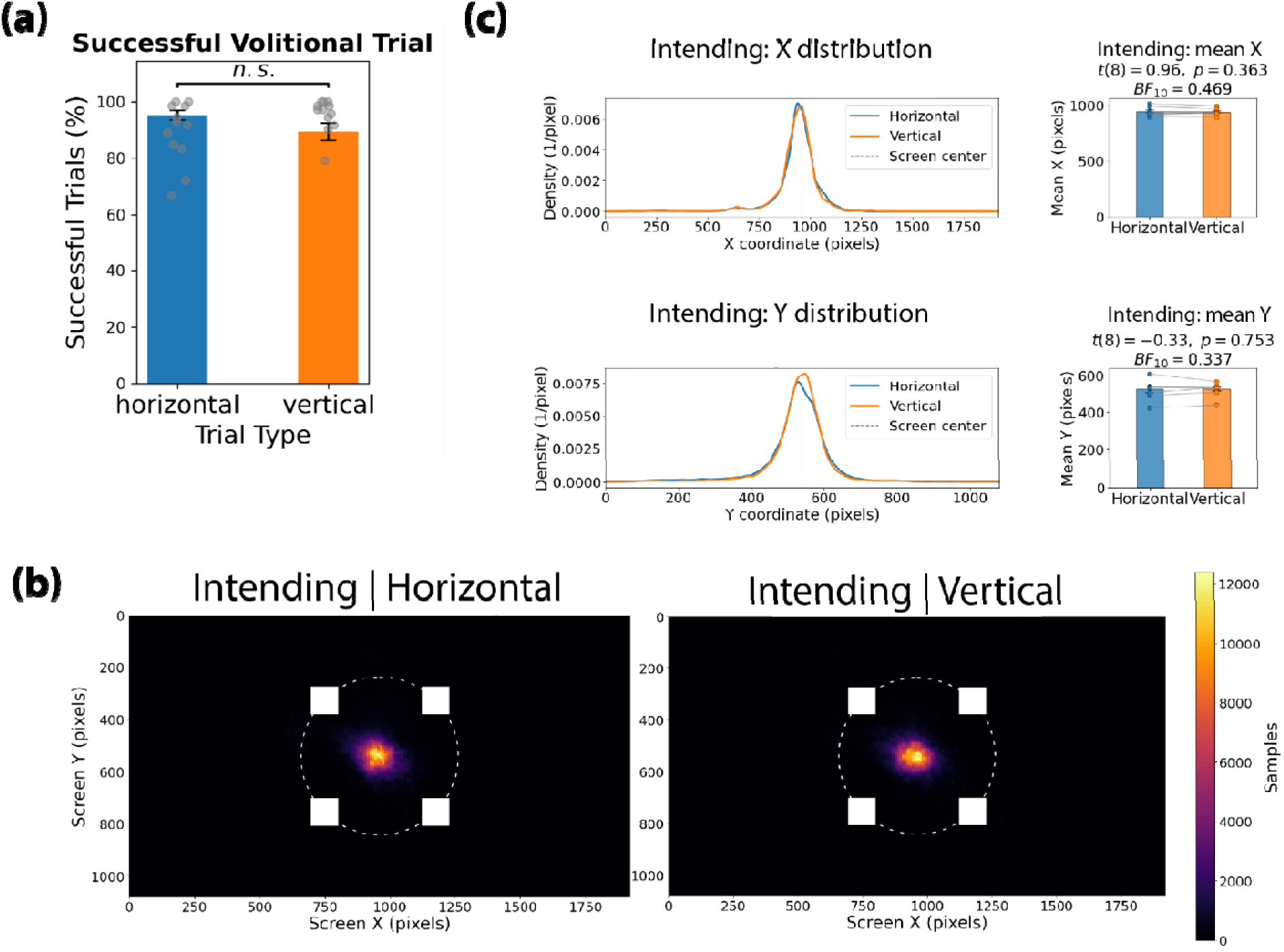
Behavioral performance and eye-position control during the intention task. (a) Percentage of successful trials for horizontal and vertical intention conditions are far above chance (50%). Points indicate individual participants; error bars indicate SEM. Success rates did not significantly differ between intention conditions (n.s.). (b) Eye-tracking heat maps during the delay period for trials in which participants intended to perceive horizontal or vertical motion, overlaid with the locations of the quartet stimuli along the aperture. The dashed circle indicates the analyzed central region. (c) Distributions of horizontal (X; top) and vertical (Y; bottom) gaze positions during the delay period for the two intention conditions. Right panels show participant-level mean gaze positions for each condition, with paired observations connected by lines.

All cross-domain decoding analyses were performed using the full set of available samples, as decoding performance was not directly compared across cross-domain conditions. Potential confounds present in the intention trials, including imbalances in left- and right-tilted square pairs or visual cue information, could not generalize to the passive viewing conditions used in the physical and ambiguous motion runs. Therefore, successful above-chance cross-domain decoding was interpreted as reflecting transfer of motion-axis representations across conditions, rather than decoding of low-level visual confounds, and no additional balancing procedures were applied.

Searchlight decoding was conducted on the *fsaverage5* surface template. Single-trial beta estimates were projected to *fsaverage5* surface space for each subject. For each cortical vertex, a local searchlight neighborhood was defined as a 10-mm-radius disc based on cortical geodesic distance computed along the cortical surface mesh using Dijkstra’s algorithm, largely following Chen et al. (2011). This approach constrained neighborhood selection to the intrinsic geometry of the cortical sheet rather than Euclidean distance in volumetric space. The *fsaverage5* surface contained 10,242 vertices per hemisphere, and an independent searchlight analysis was performed at each vertex. Within each searchlight neighborhood, beta values from all vertices in the disc were extracted and used as multivariate features for decoding horizontal versus vertical motion/intention. The same classification procedures used in the ROI analyses were independently applied at each searchlight location. Decoding accuracy from each searchlight analysis was assigned back to the center vertex, generating subject-level whole-cortex decoding maps. Group-level statistical inference was performed using a permutation-based Monte Carlo procedure. For each subject, 1,000 label permutations were used to generate subject-level null decoding maps. Group-level null distributions were then constructed by Monte Carlo sampling from the subject-level permutation maps, yielding 100,000 simulated group maps. Statistical maps were thresholded using a cluster-forming threshold of *p* < 1 × 10−5, and cluster-level significance was assessed at *p* < 1 × 10−5. This procedure effectively required both individual vertices and contiguous cluster sizes to exceed their respective null distributions derived from the permutation-based Monte Carlo simulations. Searchlight analyses were conducted using scikit-learn v1.7.1 and custom Python scripts parallelized on the Dartmouth Discovery High Performance Computing Cluster. Searchlight results were visualized using Pycortex (Gao et al., 2015) v1.3.0. Group-level statistical maps generated on the *fsaverage*5 surface were resampled to *fsaverage* resolution and displayed on flattened cortical surfaces. Additionally, we used Plotly (v.7.0.0) to project searchlight results onto the folded fsaverage5 cortical surface and visualized using an interactive three-dimensional viewer implemented (see supplementary HTML file).

## 3. Results

### 3.1 Behavioral results

The detection rate of catch trials in the intention tasks was 100%. All participants who passed pre-scan psychophysics (see Section 2.1) had a near-ceiling hit rate of perceiving motion congruent with the instructed intention in the intention task. No significant group level differences (*t* = 1.94, *p* = 0.08, *BF_10_* = 1.18) were found between vertical and horizontal trials in the intention task runs (Figure 2a). Dwell time analyses in ambiguous apparent motion task can be found in supplementary materials (Figure S8).

To assess whether differences in eye position could account for intention-related neural activity, we examined gaze behavior during the intention delay period, before the motion switch occurred. Due to eye-tracker failures during scanning, usable eye-tracking data were available for 9 of the 12 participants. Gaze-position heat maps showed that fixation remained concentrated near the center of the quartet for both horizontal and vertical intention conditions (Figure 2b). Mean horizontal (X) and vertical (Y) gaze positions did not significantly differ between intention conditions (X: *t*(8) = 0.96, *p* = 0.363, *BF10* = 0.469; Y: *t*(8) = −0.33, *p* = 0.753, *BF10* = 0.337; Figure 2c). Similar analyses of the physical and ambiguous motion runs likewise revealed no significant differences in horizontal or vertical gaze position between motion-axis conditions (Figure S1). Together, these results indicate that horizontal and vertical conditions were not associated with systematic shifts in eye position across physical perception, ambiguous perception, or intention, reducing the possibility that motion-axis-specific neural decoding was driven by differences in gaze position.

### 3.2 Univariate results

We first tested if any regions showed an overall univariate increase in activity associated with intention. We focused on visual motion-sensitive area hMT+, and then extended the analysis to the rest of cortex. To isolate activity associated with intention from visually evoked responses, we compared activity during the intention delay period with the pre-cue baseline, during which visual stimulation was identical but no motion intention had yet been specified. Significant activity was observed in hMT+ and broadly across dorsal/lateral visual and frontoparietal cortex, with comparatively limited engagement of ventral visual cortex (Figure 3b, see Figure S2 anterior display of the *t*-map).

**Figure 3.**
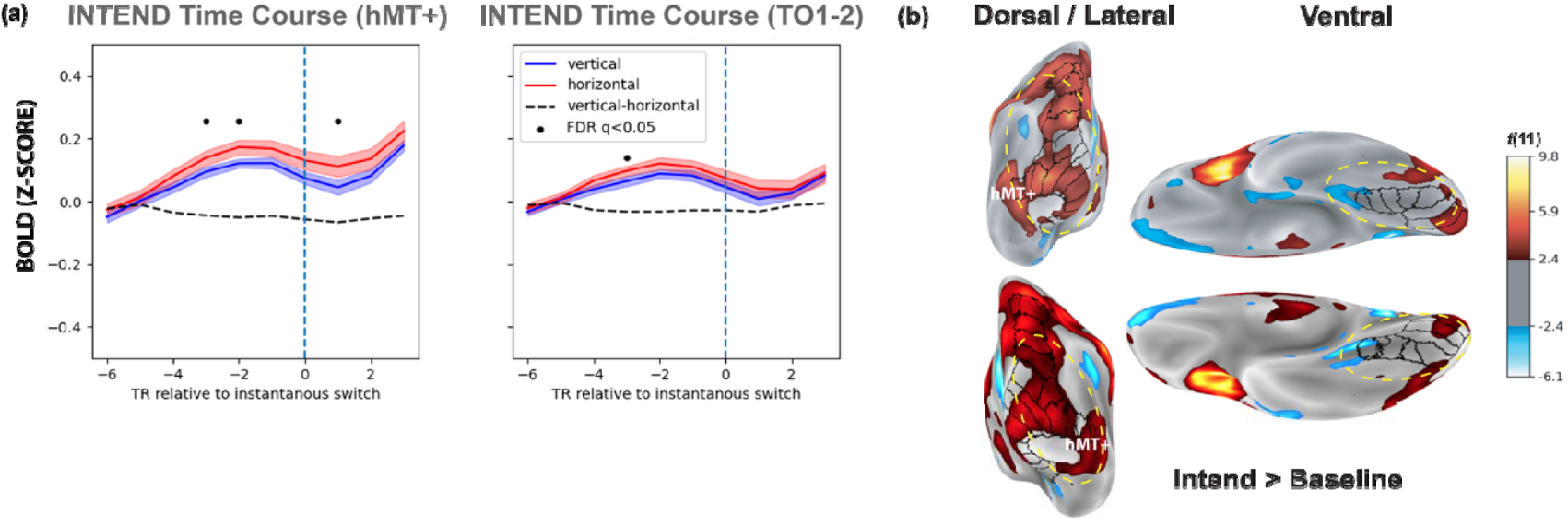
Univariate results reveal engagement of hMT+ and the broader dorsal attention network during the intention period prior to the motion stimulus (t = 0) **(a)** Univariate BOLD time-courses during the volitional intention condition in functionally defined hMT+ and atlas-defined TO1–2, aligned to the instantaneous switch (TR = 0; dashed blue line). Responses are shown separately for trials in which participants intended vertical (blue) or horizontal (red) motion; shaded regions indicate SEM across participants. The dashed black line shows the vertical-minus-horizontal difference, with black dots indicating time points surviving FDR correction (*q* < 0.05). Intention-related activity emerges during the delay period, prior to the physical switch. **(b)** Whole-brain univariate contrast of the intention period relative to baseline project to *fsaverage* template; hMT+ is indicated in white. Surface maps show group-level *t*(11) values.

Across ROIs, the univariate response increased during the intention delay and peaked before presentation of the subsequent one-shot ambiguous-motion switch (Figure 3a, see Figure S3 for time-course analysis for all Wang atlas ROIs). This temporal profile supports the interpretation that the intention-period activity modeled for subsequent decoding primarily reflects preparatory activity rather than the perceptual consequence of the switch. Additionally, at several time points and across all visual ROIs, intending to see horizontal motion consistently evoke higher BOLD amplitude compared to vertical motion.

### 3.3 hMT+ decoding results

Having established the spatial and temporal characteristics of this intention-related activity, we next asked whether its multivariate patterns contained information about the content of the intended percept. Differences in overall response magnitude between horizontal and vertical motion could potentially contribute to multivariate classification. However, such differences are not unexpected: previous fMRI work has demonstrated systematic anisotropies in cortical motion representations, including differential responses to horizontal and vertical motion across retinotopic visual cortex (Clifford et al., 2009), possibly due to disproportionate numbers of neurons turned to their respective orientations (Bauer and Dow, 1989). Thus, the univariate horizontal–vertical differences observed here may reflect an intrinsic property of motion representations rather than a condition-specific confound. Critically, if such anisotropy contributes to decoding, its generalization across physical motion, ambiguous motion perception, and volitional intention would itself provide evidence that these conditions recruit a shared representational organization for motion axis.

Prior high-resolution fMRI work (Schneider et al., 2019; Pizzuti et al., 2025) has revealed similar pseudo-columnar representations between passive viewing of physical and ambiguous apparent motion. We therefore sought to replicate these findings at standard fMRI resolution, providing a foundation for subsequent comparisons with the intention condition. In both functionally localized hMT+ and TO1–2, the equivalent retinotopically-defined areas in the Wang atlas, motion axis could be reliably decoded during passive viewing of physical and ambiguous apparent motion, with significant cross-decoding between the two conditions (all empirical *ps* < 0.001). These results indicate that physical and ambiguous motion recruit overlapping motion-axis-selective representations in hMT+ (Figure 4a).

**Figure 4.**
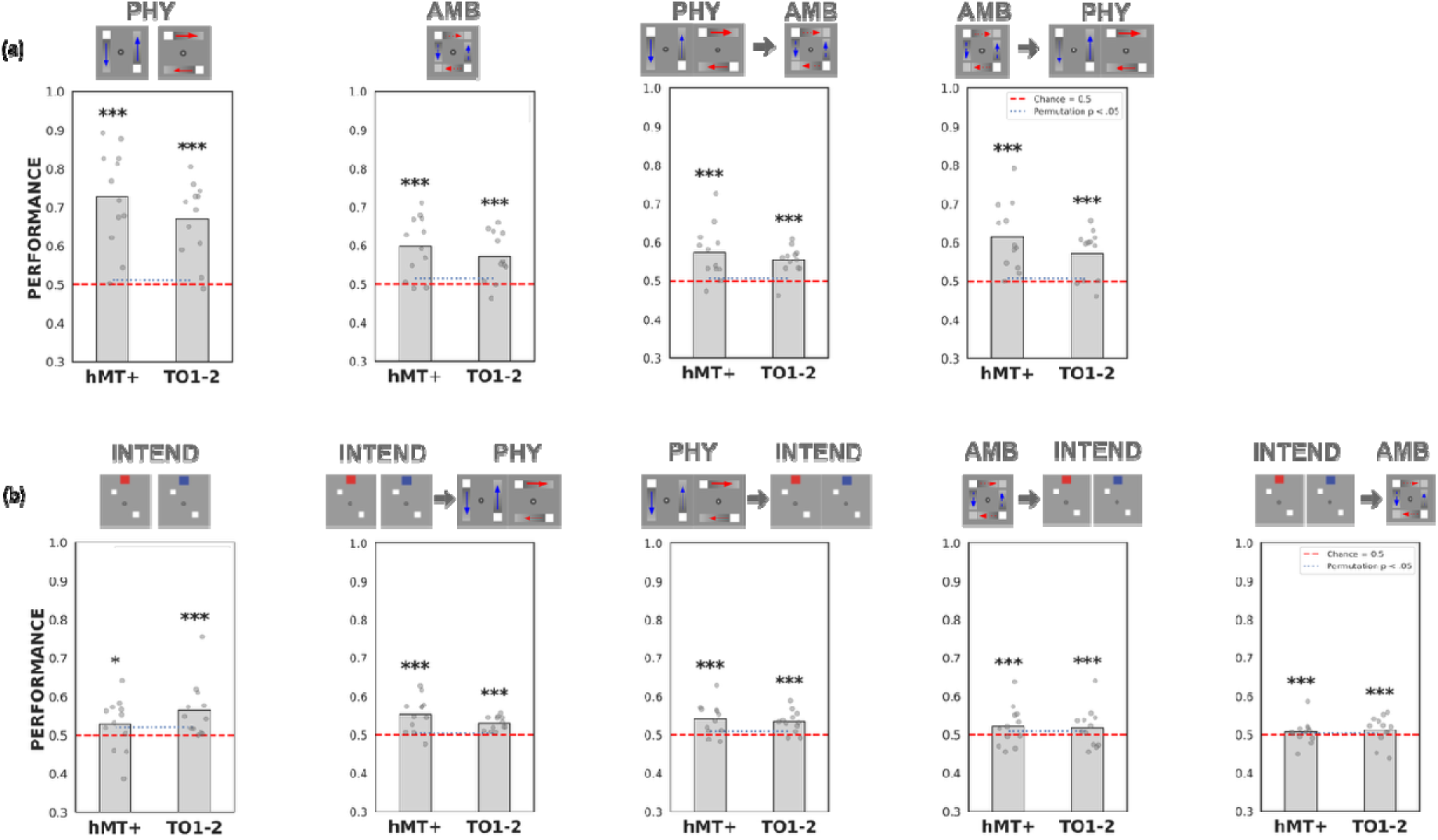
Volitional intention elicits representations similar to those evoked during passive viewing of physical and ambiguous motion in functionally defined and atlas-defined hMT+. (a) Classification and cross-classification performance (AUC-ROC) for motion axes (horizontal vs. vertical) during passive viewing of physical (PHYS) and ambiguous (AMB) motion in functionally defined hMT+ and Wang atlas-defined hMT (TO1– 2). (b) Classification and cross-classification performance within the volitional intention condition (VOL) and between VOL and passive viewing conditions. Asterisks denote statistical significance based on permutation tests (p < 0.05, p < 0.01, p < 0.001). (c) Normalized BOLD signal in TO1–2 during the intention period peaks before the one-shot switch. Within-domain decoding used leave-one-run-out cross-validation; cross-domain decoding trained on all but one run and tested on all runs of the other condition.

Critically, intention-related activity exhibited a similar representational structure (Figure 4b). Within-domain decoding of the intention delay period was comparatively weak in functionally localized hMT+ (empirical *p* < 0.05). However, cross-decoding between physical motion and intention was robust (empirical *ps* < 0.001) for both forward (train on A and test on B) and reverse (train on B and test on A) train-test pairs, as was cross-decoding between intention and ambiguous motion (empirical *ps* < 0.001 for both forward and reverse train-test pairs). Thus, despite relatively weak within-domain decoding, activity during the intention delay generalized reliably to patterns evoked during both veridical (i.e., physical) and illusory (i.e., apparent) motion perception. This suggests that intending to perceive a particular motion axis establishes a representation in hMT+ prior to the one-shot motion that has a similar neural coding structure as subsequent perceptual states. Both functional and atlas-defined hMT+ (Wang TO1–2) showed a qualitatively similar general pattern of results, despite minor differences.

### 3.4 Shared representations across perception and intention are concentrated in dorsal/lateral visual cortex

To determine whether activity recruited during intention contains motion-specific representations shared with perception, we performed cross-domain decoding between physical motion, ambiguous motion, and intention within independently defined retinotopic ROIs (Figure 5a). Cross-domain decoding revealed a striking dissociation between dorsal/lateral and ventral visual regions. Across dorsal/lateral ROIs, cross-decoding was highly robust for all condition pairs, with decoding performance significantly above chance in nearly every region (all empirical *ps* < 0.001), except in V1-V3 when training on ambiguous motion and testing on intention (empirical *p* < 0.01) and when training on physical motion and testing on ambiguous motion (empirical *p* < 0.05). In contrast, ventral visual ROIs showed little evidence for shared representations across conditions. Significant cross-decoding was observed only in VO1–2 for physical-to-intention decoding, whereas all other ventral regions failed to decode reliably above chance. To directly test this dorsal–ventral difference, we compared cross-decoding performance averaged across dorsal/lateral versus ventral ROIs within each participant (Figure 5b). Decoding performance was significantly greater in dorsal/lateral than ventral cortex for all three cross-domain comparisons (physical-to-intention, ambiguous-to-intention, and physical-to-ambiguous; all *p*s < 0.001), confirming that shared representations across perceptual and intentional states are preferentially expressed in dorsal/lateral visual cortex. Additionally, within-domain decoding in ambiguous and intention condition also shows dorsal-ventral dissociation (Figure 6c). Reversing the train–test direction in cross-decoding or using full, unbalanced samples in within-domain decoding produced a similar pattern, with strong and widespread cross-decoding throughout dorsal/lateral visual regions but only sporadic effects in ventral cortex (Figure S4, S6).

**Figure 5.**
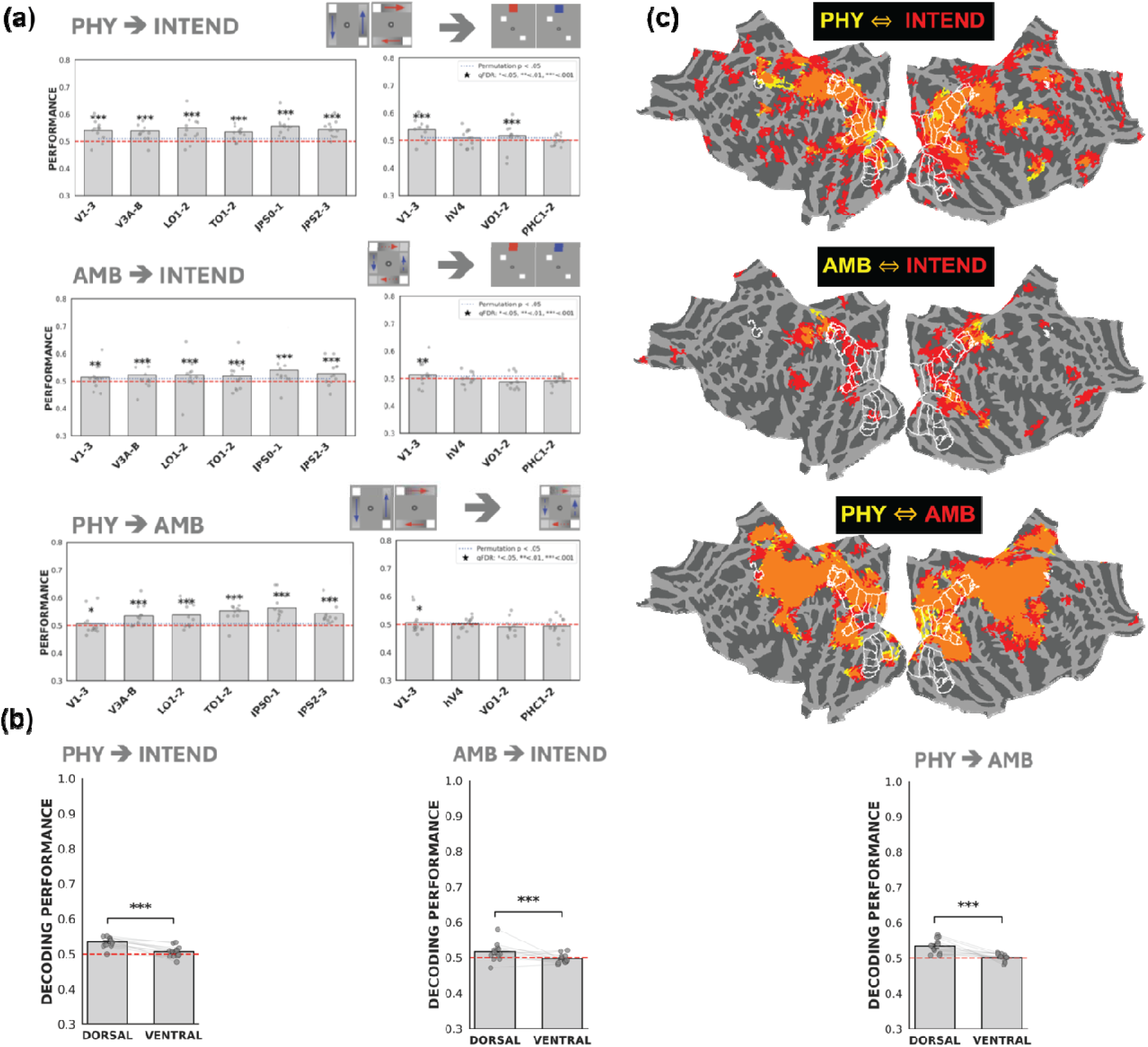
Cross-decoding between physical, ambiguous, and intended motion across dorsal/lateral and ventral visual cortex. (a) ROI-based cross-decoding performance for classifiers trained on physical motion and tested during the intention period (PHY → INTEND), trained on ambiguous motion and tested during the intention period (AMB → INTEND), and trained on physical motion and tested on ambiguous motion (PHY → AMB). Dorsal/lateral and ventral ROIs were defined using the probabilistic visual-field atlas of Wang et al. (2015). Bars indicate mean decoding performance across participants; individual dots indicate participant-level values. Asterisks indicate statistical significance based on permutation testing with FDR correction where applicable. Results for the corresponding reverse train–test directions are reported in the Supplementary Material. (b) Comparisons between dorsal and ventral cortex cross-decoding performance. Significance was assessed using permutation tests with Benjamini–Hochberg FDR correction; (c) Surface-based searchlight cross-decoding maps for the same representational comparisons. Yellow indicates vertices showing significant cross-decoding in the forward training-to-test direction (PHY → INTEND, AMB → INTEND, or PHY → AMB), red indicates significant cross-decoding in the corresponding reverse direction (INTEND → PHY, INTEND → AMB, or AMB → PHY), and orange indicates spatial overlap between significant forward and reverse cross-decoding. White contours indicate atlas-defined visual ROIs. Cluster forming threshold of *p*<1e-05 was applied; final group level maps were then clustered at *p*<1e-05 using a group level null distribution generated using bootstrapping individual null maps with replacement (see Methods section 2.6). Significance was assessed using permutation tests with Benjamini–Hochberg FDR correction; *q* < 0.05 (*), *q* < 0.01 (**), and *q* < 0.001 (***).

**Figure 6.**
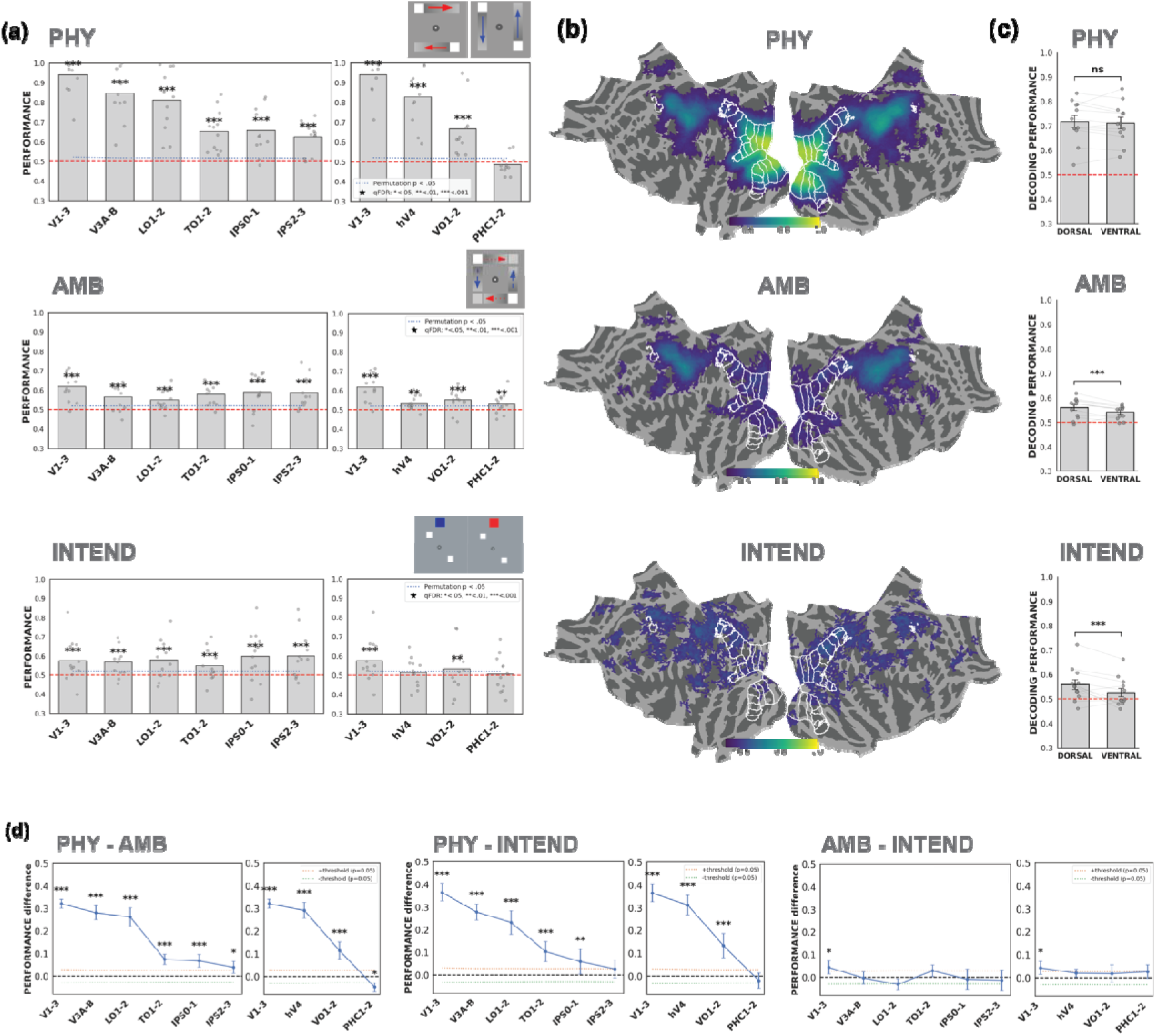
Representational strength of veridical perception, illusion, and intention in dorsal/lateral and ventral visual cortex. (a) ROI-based representational strength. Decoding performance (AUC-ROC) for motion axis (horizontal vs. vertical) using leave-one-run-out cross-validation, indexing representational strength within each condition: physical (PHY), ambiguous (AMB), and intention. Bars indicate group-level mean decoding performance across participants, with individual participant values overlaid. The blue dotted line indicates the *p* = 0.05 threshold derived from group-level permutation tests. Asterisks denote significance based on permutation testing with FDR correction where applicable. (b) Pairwise comparisons of representational strength. Differences in decoding performance between conditions (PHY – AMB, PHY – INTEND, and AMB – INTEND). Points indicate mean differences across participants, with error bars indicating SEM. Positive values indicate stronger representations in the first condition of each contrast. Horizontal dashed lines indicate permutation-derived significance thresholds; asterisks denote statistical significance with FDR correction where applicable. (c) (d) Comparisons between dorsal and ventral cortex cross-decoding performance. (d) Searchlight decoding performance for motion axis (horizontal vs. vertical) within the physical, ambiguous, and intend conditions. Surface color indicates local cross-validated decoding performance, with higher values reflecting stronger local information about the represented motion axis. White contours indicate atlas-defined visual ROIs. Cluster forming threshold of p<1e-05 was applied; final group level maps were then clustered at p<1e-05 using a group level null distribution generated using bootstrapping individual null maps with replacement (see Methods section 2.6). Significance was assessed using permutation tests with Benjamini–Hochberg FDR correction; *q* < 0.05 (*), *q* < 0.01 (**), and *q* < 0.001 (***).

Our ROIs were defined using a probabilistic atlas rather than subject-specific functional localization, limiting the anatomical precision of EVC measurements. Consequently, we were unable to replicate previous reports that apparent motion evokes BOLD activity along the perceived motion path in V1 (Muckli et al., 2005), as no subject specific localizer data on motion path was collected. This methodological limitation may also have contributed to the relatively weak within-domain decoding and cross-decoding between apparent motion and intention observed in EVC. Therefore, although our results should not be taken as evidence that EVC plays only a limited role in rendering vivid perception of apparent motion or representing intention to see apparent motion.

### 3.5 Representational strength analysis distinguishes ambiguous motion and intention from physical motion

Having established that physical motion, ambiguous motion, and intention share representational patterns within dorsal/lateral visual cortex, we next asked whether these representations were expressed with comparable strength across conditions. We quantified representational strength using within-domain motion-axis decoding with leave-one-run-out (LOO) cross-validation with trial balanced (see Methods section 2.6) data (Figure 6a). For physical motion, decoding was significantly above chance in all dorsal/lateral ROIs (V1–3, V3A–B, LO1–2, TO1–2, IPS0–1, and IPS2–3; all empirical *ps* < 0.001) and in V1–3, hV4, and VO1–2 in ventral cortex (all empirical *ps* < 0.001), but not PHC1–2. For ambiguous motion, decoding was significant throughout dorsal/lateral cortex (all empirical *ps* < 0.001) and across ventral ROIs (V1–3 and VO1–2, empirical *ps* < 0.001; hV4 and PHC1–2, empirical *ps* < 0.01). For intention, decoding was similarly significant throughout dorsal/lateral cortex (all empirical *ps* < 0.001), whereas ventral effects were restricted to V1–3 (empirical *p* < 0.001) and VO1–2 (empirical *p* < 0.01), with no significant decoding in hV4 or PHC1–2. All reported significant effects survived FDR correction across ROIs and train-test pairs. Trial balanced results do not differ qualitatively with full sample decoding results (Figure S5a).

We then compared within-domain decoding performance between the three conditions within each ROI (Figure 6d). Physical motion produced substantially stronger and more reliable motion-axis representations than either ambiguous motion or intention, particularly in early visual and dorsal/lateral regions. Pairwise comparisons (see Methods section 2.6) confirmed significantly greater decoding performance for physical relative to ambiguous motion (PHY – AMB): empirical *ps* < 0.001 in V1–3, V3A/B, LO1/2, TO1/2, and IPS0/1; *p* < 0.05 in IPS2/3. Similarly, decoding was greater for physical motion relative to intention (PHY – INTEND): empirical *ps* < 0.001 in V1–3, V3A/B, LO1/2, TO1/2, and hV4; *p* < 0.01 in IPS0/1. In contrast, decoding performance did not reliably differ between ambiguous motion and intention (AMB - INTEND) in either dorsal/lateral or ventral ROIs, except in V1–3 (*p* < 0.05), with no other comparisons surviving FDR correction. Analyses of the full, unbalanced dataset yielded qualitatively similar results (Figure S5b), indicating that the observed pattern was robust to trial balancing.

Interestingly, within-domain decoding also revealed a similar dorsal–ventral organization for ambiguous motion and intention (Figure 6c). Decoding performance was significantly greater in dorsal/lateral than ventral visual cortex for both ambiguous motion and intention (both empirical *p*s < 0.001), whereas no dorsal–ventral difference was observed for physical motion (empirical *p* > 0.05). Thus, although ambiguous motion and intention differ in their source, one reflecting perceptual resolution of an ambiguous stimulus and the other an internally specified perceptual goal, both showed a similar representational strength across retinotopic cortex and a comparable dorsal–ventral bias that differs from unambiguous physical motion. Using unbalanced full sample data revealed similar patterns (Figure S6b). Together, these findings suggest that intention establishes a prospective motion representation resembling the neural state associated with illusory motion perception.

Complete statistics for all ROI decoding analyses are reported in supplementary table 1–7.

### 3.6 Entire-Cortex Searchlight Reveals Additional shared Frontal Representations

To determine whether this dorsal/lateral organization extended beyond the predefined ROIs, we complemented the ROI analyses with entire-cortex searchlight decoding (Figure 5c, 6b). Consistent with cross-decoding ROI results, significant cross-decoding between physical motion and intention, as well as between ambiguous motion and intention, was concentrated within dorsal/lateral visual cortex and closely followed the distribution of retinotopically defined dorsal/lateral visual regions (Figure 5c). For entire-cortex searchlight within-domain decoding analysis, consistent with corresponding ROI results, physical motion exhibited a pronounced posterior-to-anterior gradient, with classification strongest in early visual cortex and progressively weaker in downstream regions. In contrast, ambiguous motion and volitional intention exhibited markedly flatter representational profiles across the visual hierarchy. Motion-axis information was broadly decodable throughout dorsal/lateral visual regions during ambiguous motion and remained reliably decodable in dorsal/lateral cortex during the intention period (Figure 6b).

Beyond retinotopic occipital and parietal cortex, significant searchlight clusters were also observed in frontal regions, including the superior precentral sulcus (sPCS) near the Wang atlas FEF boundary, a region implicated in the frontoparietal circuitry supporting top-down attentional control (Kastner and Ungerleider, 2000). These clusters were present in physical-to-intention cross-decoding and within-domain decoding for all three conditions, although they did not survive the cluster threshold for ambiguous-to-intention cross-decoding. Additional frontal clusters were observed in searchlight and univariate analyses, particularly for within-domain decoding of intention (see supplementary interactive view of within and cross-domain decoding results), in regions around dACC, DLPFC, and possibly the insula. These regions may correspond to portions of the fronto-parietal and cingulo-opercular networks described by Dosenbach et al (2008). It is important to note, however, that frontal association cortex exhibits substantial inter-individual variability in its functional organization, and individual-specific functional boundaries can diverge considerably from group-average organization (Mueller et al., 2013; Gordon et al., 2017), making precise anatomical attribution of group-level effects within frontal cortex less certain, particularly given our relatively limited sample size (N = 12). We therefore interpret these effects conservatively as evidence that intention engages a broader frontoparietal network.

In the physical and ambiguous motion conditions, horizontal and vertical reports were made with different hands, introducing a potential motor confound that could contribute to decodable activity along the sensorimotor cortex, while significant searchlight clusters during the intention period were largely absent from primary sensorimotor cortex and the central sulcus (Figure 5c, 6b), due to our design to prevent motor-preparatory confound during intention (see Discussion).

## 4. Discussion

The present study investigated whether a top-down intention (to perceive an upcoming ambiguous motion stimulus as moving horizontally or vertically) is sufficient to alter sensory representations prior to the bottom-up stimulus change that results in a conscious perception of motion corresponding to the intended motion. Using a cue–delay paradigm designed to isolate intention-related activity from the perceptual rivalry inherent to the apparent motion quartet, we found that intending to perceive a particular motion axis evokes neural representations resembling those associated with both veridical (i.e., physical) and illusory (i.e., apparent) motion perception. Across functionally localized hMT+ (Figure 4), dorsal/lateral retinotopic visual regions (Figure 5a, 6a), and whole-cortex searchlight analyses (Figure 5c, 6b), intention-related activity generalized to perceptual and illusory motion states through robust cross-decoding. These findings suggest that an observer’s intention can establish prospective sensory representations prior to stimulus presentation (i.e., prior to the dot pair switch) and that these representations share a common neural code with subsequent perceptual experience. Temporal analyses further showed that activity peaked before the onset of the motion switch, confirming that the decoded signals primarily reflected intention-related processes rather than responses to the subsequent sensory event (Figure 3a; Figure S3). Our findings extend previous work demonstrating motion-axis-selective representations in hMT+ during physical and ambiguous apparent motion perception (Muckli et al., 2002; Schneider et al., 2019), suggesting that intention recruits neural populations that are also engaged during motion perception.

Intention-related representations were preferentially expressed in dorsal/lateral visual regions, with robust cross-decoding between intention and both physical and ambiguous motion in V3AB, hMT+, IPS, and sPCS regions, but substantially weaker cross-decoding in ventral visual cortex (Figure 5; Figure 6). This pattern emerged consistently across ROI analyses and whole-cortex searchlight analyses and therefore does not depend on a particular ROI definition strategy. Moreover, regions exhibiting successful cross-decoding overlapped substantially with regions showing increased univariate activity during the intention period relative to baseline (Figure 3b, 5c, 6b). Because many of these regions overlap with the dorsal attention network, one possibility is that intention-related representations are supported by top-down feedback originating from attentional control systems. Such feedback could bias motion-sensitive populations toward the intended percept in a manner analogous to feature- or spatial-attention (Reynolds et al., 2000; Treue & Martínez-Trujillo, 1999). Although our data cannot establish causality, they are consistent with higher-order control regions shaping subsequent perceptual representations.

Intention-period decoding is unlikely to reflect motor preparation: participants did not know which motor response would be required until after the intention period ended, and significant searchlight clusters during this period were largely absent from sensorimotor cortex. Although motor activity may contribute to decoding in the physical and ambiguous motion conditions, it cannot account for cross-decoding with the intention condition, supporting the interpretation that generalization reflects shared perceptual and intention-related representations.

Preparatory sensory representations have previously been demonstrated in human fMRI studies of attentional templates, in which representations of anticipated targets, or associated information useful for prioritizing them, can be detected in sensory and prefrontal cortex before stimulus onset, which may facilitate subsequent attentional selection (Stokes et al., 2009; Zhou and Geng, 2025). Attentional templates prepare the visual system to select task-relevant sensory information, whereas our paradigm requires participants to voluntarily bias one of two competing interpretations of identical ambiguous input. Our study extends this work by showing that preparatory representation reflects not only what observers expect to detect, but what they intend to see.

Representational strength analyses further revealed an important distinction between physical motion perception and the other two conditions. Physical motion produced substantially stronger within-domain decoding performance than either ambiguous motion or intention, particularly within the early visual cortex (Figure 6a,d; Figure S5). This effect was also evident in the searchlight analyses (Figure 6b), where physical motion exhibited a pronounced posterior-to-anterior gradient, with the strongest decoding concentrated in early visual cortex, with progressively weaker representations in more anterior regions. This profile is consistent with a representation dominated by feedforward sensory input, represented most strongly at the earliest stages of cortical processing and then gradually transformed through subsequent stages of the visual hierarchy (Felleman and Van Essen, 1991). In contrast, ambiguous motion and intention displayed remarkably similar representational profiles. Decoding performance was statistically indistinguishable between the two conditions across nearly all ROIs, and both exhibited a relatively stable distribution of decoding strength across dorsal/lateral visual regions rather than exhibiting strong early-visual dominance. This similarity is particularly striking because the two conditions differ fundamentally in their sensory input. During ambiguous motion, perceptual representations emerge through the resolution of competing interpretations generated by alternating motion inducers during passive viewing. During the intention period, however, no motion stimulus was present. Nevertheless, endogenous intention generated neural representations that were statistically indistinguishable from those elicited by the illusory percept itself and generalized robustly across conditions through cross-decoding. A plausible interpretation of these findings is that physical motion perception is characterized primarily by feedforward sensory processing, whereas ambiguous motion perception and intention both rely more heavily on top-down influences distributed throughout the dorsal/lateral visual hierarchy. Under this account, the similarity between intention and ambiguous motion arises because both involve the construction of a motion representation rather than the direct encoding of a physically present motion signal. It follows that the prospective sensory state established by intention more closely resembles the neural representation generated during the perceptual resolution of ambiguity than the one generated by physical motion.

An important question concerns the precise cognitive mechanisms responsible for the observed preparatory representations. Participants may have attended selectively to a motion-axis feature as in classic feature-based attention tasks (Serences and Boynton, 2007; Silver et al., 2007), imagined a particular motion trajectory in anticipation of the stimulus (Dijkstra et al., 2017; Dijkstra et al., 2019; Li et al., 2023), actively maintained an intended perceptual outcome (Albers et al., 2013), or increased their expectation to a specific motion event (Kok et al., 2017; de Lange et al., 2018) — and these processes are likely not mutually exclusive. Indeed, intention, attention, expectation, and imagery often co-activate the frontal-parietal network in paradigms like ours, and it may not be possible to fully dissociate their contributions (Nau et al., 2024). Moreover, the influence of intention on sensory cortex may be implemented through mechanisms similar to those engaged by attention, such as feature-selective gain modulation that enhances neural populations representing the intended motion axis. Thus, intention and attention need not constitute mutually exclusive explanations: intention may specify the desired perceptual outcome while recruiting attentional mechanisms to bias sensory processing of subsequent bottom-up input toward the intended outcome.

Crucially, whatever cognitive process participants used to implement their intention, it produced motion-axis-specific sensory representations — similar in strength and cortical organization from those observed during ambiguous motion perception itself. This representational similarity to perception, rather than to a neutral preparatory state, is the key finding: the brain did not merely prepare for motion in general, but pre-configured sensory cortex in a way that resembled seeing it.

Future studies employing high-resolution laminar fMRI may help clarify the circuit-level basis of these top-down influences in mesoscopic-scale (e.g. Bergmann et al., 2024). Because feedforward and feedback signals preferentially target different cortical layers (Stephan et al., 2019), laminar measurements could determine whether intention-related activity reflects feedback arriving from higher-order cortical regions and help identify potential sources of these signals present in the sensory cortex.

We conclude that the mere intention to perceive a stimulus in a particular way is sufficient to establish motion-axis-specific representations in early sensory cortex – prior to any stimulus change and prior to conscious perceptual experience. Critically, these preparatory activity patterns closely resemble those observed during actual perception of both physical and illusory motion, suggesting that intention pre-configures sensory cortex in a perception-like state. Overall, our findings demonstrate that volitional top-down control can shape early sensory cortex prior to perceptual experience. As these areas are core nodes of the dorsal attentional network, it is likely that endogenous attention and intention may rely on a shared neural mechanism, such as gain control.

## Acknowledgements

We thank Ahmad Wahab and Adeola Afonja for assistance with data collection, Marvin Maechler for providing customed code to visualize brain maps, Caroline Robertson for comments and suggestions throughout various stages of the project, and the graduate students and postdoctoral researchers who provided valuable feedback during the Vision Science Joint Lab meetings at Dartmouth department of Psychological and Brain Sciences.

## Author Contributions

Conceptualization: W.S., E.C., N.H.H., P.C., P.U.T.; Methodology: W.S., E.C., P.J.K.; Data curation: W.S., E.C., N.H.H.; Formal analysis: W.S., E.C.; Software: W.S., E.C., P.J.K.; Writing – original draft: W.S.; Writing-review & editing: E.C., V.S.S., P.U.T., P.C., P.J.K.; Supervision: P.U.T., N.H.H., P.C., V.S.S., P.J.K.; Funding acquisition: P.U.T., N.H.H., P.C., V.S.S.

## Data and Code Availability

The fMRI data used in this study are available through OpenNeuro at openneuro.org/datasets/ds008704. Behavioral data, stimuli code and custom analysis code, and interactive searchlight results html file were uploaded to OSF: https://osf.io/6smxf.

