## Supplementary for "Seeing at Will: Shared Neural Representations of Motion Perception and Intention"

***Text S1:***

**Pre-scan Psychophysics details**

Apparatus and stimuli

Prior to fMRI scanning, participants completed a psychophysical session to determine an individualized stimulus configuration for the ambiguous motion quartet and to assess their ability to volitionally bias its perceived motion direction. Stimulus presentation and response collection were controlled using PsychoPy.

The apparent-motion quartet consisted of four 1° × 1° white squares on a grey background positioned at the corners of an invisible rectangle centered on fixation. On each stimulus frame, two diagonally opposed squares were presented simultaneously and alternated with the other diagonal pair, producing apparent motion that could be perceived as either horizontal or vertical. The quartet was positioned within a circular aperture with a radius of 3° of visual angle. A central fixation point (radius = 0.1°) remained present throughout the experiment, and participants were instructed to maintain fixation. Stimuli were presented on a 60-Hz display (1920 × 1080 pixels; approximately 52.3 × 29.4 cm) at a viewing distance of 68 cm.

The relative horizontal and vertical spacing of the quartet was manipulated by varying its aspect ratio while maintaining a constant overall eccentricity, so that all four quartet squares were equidistant from fixation. Changes in aspect ratio systematically altered the relative tendency to perceive horizontal versus vertical motion. Because the aspect ratio producing approximately balanced horizontal and vertical perception varies across observers, an individualized aspect ratio was determined for each participant.

The pre-scan session consisted of three successive stages: (1) an initial estimation using the method of limits, (2) a more precise estimation using the method of constant stimuli, and (3) an assessment of volitional control.

Method of limits

The first stage provided an initial estimate of the aspect ratio at which perception transitioned between horizontal and vertical apparent motion. Participants completed one practice trial followed by two runs of 10 trials each, for a total of 20 experimental trials.

During each trial, the two diagonal configurations of the quartet alternated every 250 ms while the aspect ratio changed progressively. On half of the trials, the aspect ratio progressed from a configuration favoring one motion direction toward a configuration favoring the other (ascending trials), whereas on the remaining trials the progression was reversed (descending trials). Ascending and descending trials were intermixed within each run. The initial diagonal pair was also counterbalanced across trials.

Participants were instructed to maintain fixation and press the space bar as soon as they experienced a change in the perceived direction of apparent motion. The aspect ratio at the time of the response was recorded as the perceptual transition point. Following the response, the fixation point turned green for 1-s to provide feedback, followed by a 1-s intertrial interval.

Transition points were estimated separately for ascending and descending trials and then combined to obtain an initial estimate of the aspect ratio at which horizontal and vertical apparent-motion percepts were approximately balanced. This estimate was used to center the range of aspect ratios presented during the subsequent method-of-constant-stimuli procedure.

Method of constant stimuli

The second stage provided a more precise estimate of each participant’s point of subjective equality (PSE). Eight aspect ratios centered around the transition estimate obtained from the method of limits were selected individually for each participant.

Participants completed four runs of 80 trials each, yielding 320 trials in total. Each of the eight aspect ratios was presented 10 times per run and therefore 40 times across the entire procedure. Trial order was randomized within each run, and the diagonal pair presented first was counterbalanced across trials.

On each trial, the first diagonal pair of the quartet was presented for 500 ms, followed by the alternate diagonal pair for 500 ms, resulting in a 1-s apparent-motion presentation. Participants then had up to 5-s to report whether they had perceived horizontal or vertical motion. They were instructed to report the motion direction they actually perceived without attempting to intentionally influence their percept during this stage.

Following each response, the fixation point turned green for 1-s, followed by a 1-s intertrial interval. Responses were collected separately for each aspect ratio, and a psychometric function was fit to the proportion of horizontal and vertical reports. The aspect ratio corresponding to equal probabilities of perceiving horizontal and vertical motion was defined as the participant’s PSE.

The individually estimated PSE was subsequently used as the ambiguous quartet configuration for the volitional-control screening and the fMRI experiment. This procedure minimized pre-existing stimulus-driven biases toward either horizontal or vertical motion and provided a stimulus that was approximately perceptually balanced for each participant.

Pre-scan assessment of volitional control

The final stage assessed participants’ ability to intentionally bias their perception of the ambiguous motion quartet. The quartet was presented at the participant-specific PSE determined in the preceding procedure, such that the physical spatial configuration of the quartet was identical across horizontal- and vertical-intention trials.

Participants completed four runs of 40 trials each, yielding 160 trials in total, equal to the total number of volitional trials subsequently presented during fMRI scanning. Horizontal and vertical intention trials occurred equally often and were randomized within each run. The initial diagonal configuration of the quartet was independently counterbalanced across trials.

At the beginning of each trial, participants were instructed to try to perceive the quartet as moving either horizontally or vertically. The intention cue was presented for 1-s together with the initial configuration of the quartet. Following the cue, the quartet was presented at the participant’s individualized PSE. The first diagonal configuration remained visible for 4-s before alternating to the other diagonal configuration for 500 ms, producing the critical apparent-motion event.

After stimulus presentation, participants had up to 5-s to report the motion direction they had actually perceived. Participants were explicitly instructed to report their percept rather than simply report the instructed direction. Following the response, the fixation point turned green for 1 s, followed by a 1-s intertrial interval.

A trial was classified as successful when the reported motion direction matched the instructed direction. Volitional-control performance was calculated as the proportion of successful trials across horizontal and vertical intention conditions.

Prior to scanning, all participants except the authors (N.H., E.C., and W.S.) were screened for volitional control using this procedure. Participants with an overall hit rate greater than 70% were eligible to proceed to scanning. One participant did not meet this criterion and was excluded prior to scanning.

***Text S2*:**

**MRI preprocessing details using fMRIPrep 24.1.1**

A total of 2 fieldmaps were found within the input BIDS structure for each subject (one in each session). A *B0*-nonuniformity map (or *fieldmap*) was estimated based on two echo-planar imaging (EPI) references with top-up (Andersson et al., 2003) for each session based on the fieldmap collected in the beginning of the session.

One T1-weighted (T1w) image was extracted from the input BIDS dataset. The T1w image was corrected for intensity non-uniformity (INU) with *N4BiasFieldCorrection* (Tustison et al. 2010), distributed with ANTs 2.5.3 (Avants et al. 2008, RRID:SCR_004757), and used as T1w-reference throughout the workflow. The T1w-reference was then skull-stripped with a *Nipype* implementation of the *antsBrainExtraction.sh* workflow (from ANTs), using OASIS30ANTs as target template. Brain tissue segmentation of cerebrospinal fluid (CSF), white-matter (WM) and gray-matter (GM) was performed on the brain-extracted T1w using *fast* (FSL (version 6.0.7.11), RRID:SCR_002823, Zhang et al., 2001). Brain surfaces were reconstructed using *recon-all* (FreeSurfer 7.3.2, RRID:SCR_001847, Dale, Fischl, and Sereno 1999), and the brain mask estimated previously was refined with a custom variation of the method to reconcile ANTs-derived and FreeSurfer-derived segmentations of the cortical gray-matter of Mindboggle (RRID:SCR_002438, Klein et al. 2017). Volume-based spatial normalization to one standard space (MNI152NLin2009cAsym) was performed through nonlinear registration with *antsRegistration* (ANTs 2.5.3), using brain-extracted versions of both T1w reference and the T1w template. The following template was selected for spatial normalization and accessed with *TemplateFlow* (24.2.0, Ciric et al. 2022): *ICBM 152 Nonlinear Asymmetrical template version 2009c* [Fonov et al. (2009), RRID:SCR_008796; TemplateFlow ID: MNI152NLin2009cAsym].

For each of the 31 BOLD runs found per subject (across all tasks and sessions), the following preprocessing was performed. First, a reference volume was generated, using a custom methodology of *fMRIPrep*, for use in head motion correction. Head-motion parameters with respect to the BOLD reference (transformation matrices, and six corresponding rotation and translation parameters) are estimated before any spatiotemporal filtering using *mcflirt* (FSL, Jenkinson et al. 2002). The estimated *fieldmap* was then aligned with rigid-registration to the target EPI (echo-planar imaging) reference run. The field coefficients were mapped on to the reference EPI using the transform. The BOLD reference was then co-registered to the T1w reference using *bbregister* (FreeSurfer) which implements boundary-based registration (Greve and Fischl 2009). Co-registration was configured with six degrees of freedom. Several confounding time-series were calculated based on the preprocessed BOLD: framewise displacement (FD), DVARS and three region-wise global signals. FD was computed using two formulations following Power (absolute sum of relative motions, Power et al. (2014)) and Jenkinson (relative root mean square displacement between affines, Jenkinson et al. (2002)). FD and DVARS are calculated for each functional run, both using their implementations in *Nipype* (following the definitions by Power et al. 2014). The three global signals are extracted within the cerebrospinal fluid (CSF), the white matter (WM), and the whole-brain masks. Additionally, a set of physiological regressors were extracted to allow for component-based noise correction (*CompCor*, Behzadi et al. 2007). Principal components are estimated after high-pass filtering the *preprocessed BOLD* time-series (using a discrete cosine filter with 128s cut-off) for the two *CompCor* variants: temporal (tCompCor) and anatomical (aCompCor). tCompCor components are then calculated from the top 2% variable voxels within the brain mask. For aCompCor, three probabilistic masks (CSF, WM and combined CSF+WM) are generated in anatomical space. The implementation differs from that of Behzadi et al. (2007) in that instead of eroding the masks by 2 pixels on BOLD space, a mask of pixels that likely contain a volume fraction of gray matter (GM) is subtracted from the aCompCor masks. This mask is obtained by dilating a GM mask extracted from the FreeSurfer’s *aseg* segmentation, and it ensures components are not extracted from voxels containing a minimal fraction of GM. Finally, these masks are resampled into BOLD space and binarized by thresholding at 0.99 (as in the original implementation). Components are also calculated separately within the WM and CSF masks. For each CompCor decomposition, the *k* components with the largest singular values are retained, such that the retained components’ time series are sufficient to explain 50 percent of variance across the nuisance mask (CSF, WM, combined, or temporal). The remaining components are dropped from consideration. The head-motion estimates calculated in the correction step were also placed within the corresponding confounds file. The confound time series derived from head motion estimates and global signals were expanded with the inclusion of temporal derivatives and quadratic terms for each (Satterthwaite et al. 2013). Frames that exceeded a threshold of 0.5 mm FD or 1.5 standardized DVARS were annotated as motion outliers. Additional nuisance time series are calculated by means of principal components analysis of the signal found within a thin band (*crown*) of voxels around the edge of the brain, as proposed by (Patriat, Reynolds, and Birn 2017). The BOLD time-series were resampled onto the following surfaces (FreeSurfer reconstruction nomenclature): *fsnative*, *fsaverage*, and downsampled *fsaverage5* for the purpose of conducting whole-cortex searchlight analyses. All re-samplings can be performed with *a single interpolation step* by composing all the pertinent transformations (i.e. head-motion transform matrices, susceptibility distortion correction when available, and co-registrations to anatomical and output spaces). Gridded (volumetric) re-samplings were performed using *nitransforms*, configured with cubic B-spline interpolation. Non-gridded (surface) re-samplings were performed using *mri_vol2surf* (FreeSurfer).

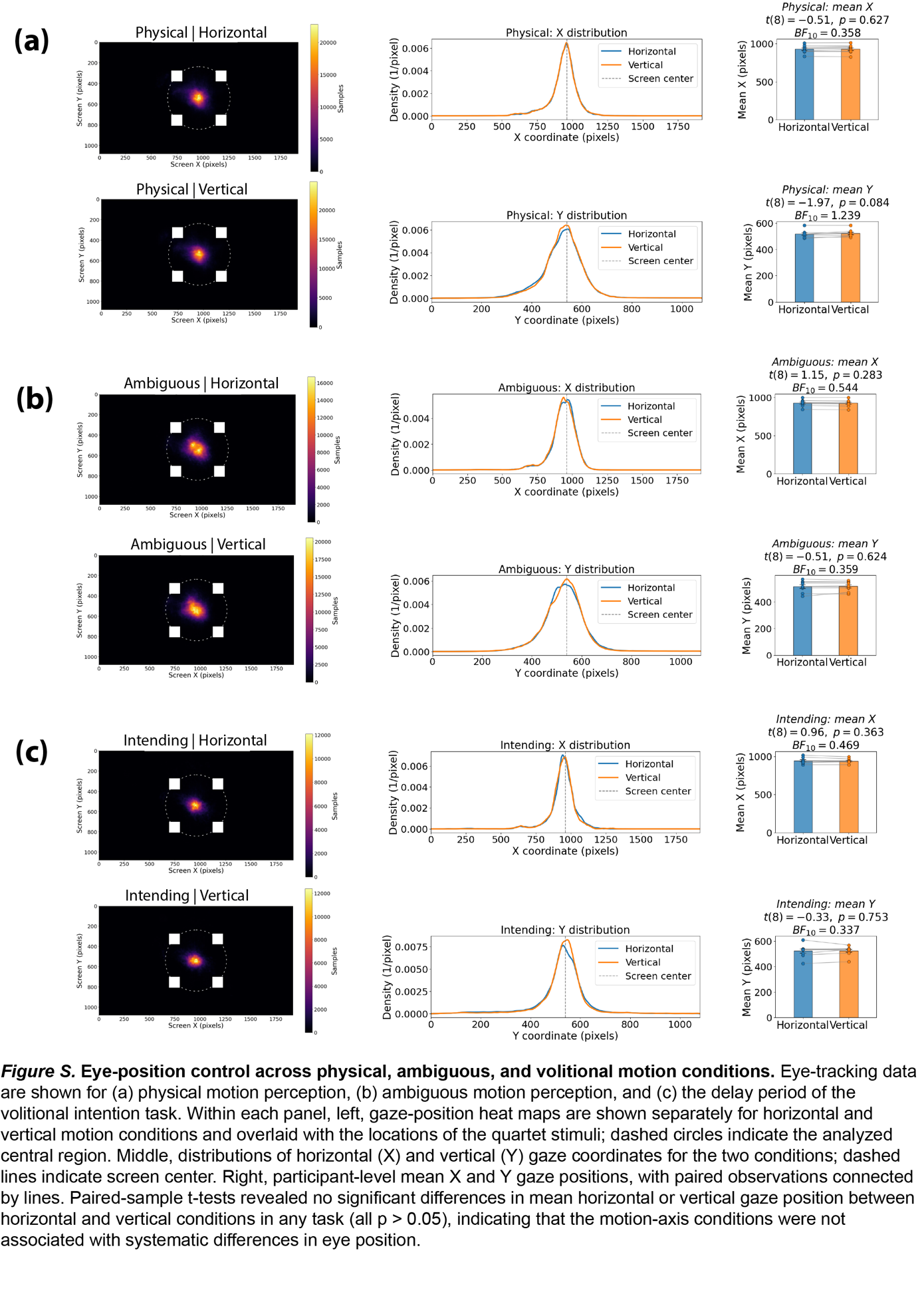

***Figure S1.* Eye-position control across physical, ambiguous motion runs and volitional intention runs.** Eye-tracking data are shown for (a) physical motion perception, (b) ambiguous motion perception, and (c) the delay period of the volitional intention task. Within each panel, left, gaze-position heat maps are shown separately for horizontal and vertical motion conditions and overlaid with the locations of the quartet stimuli; dashed circles indicate the analyzed central region. Middle, distributions of horizontal (X) and vertical (Y) gaze coordinates for the two conditions; dashed lines indicate screen center. Right, participant-level mean X and Y gaze positions, with paired observations connected by lines. Paired-sample t-tests revealed no significant differences in mean horizontal or vertical gaze position between horizontal and vertical conditions in any task (all p > 0.05), indicating that the motion-axis conditions were not associated with systematic differences in eye position.

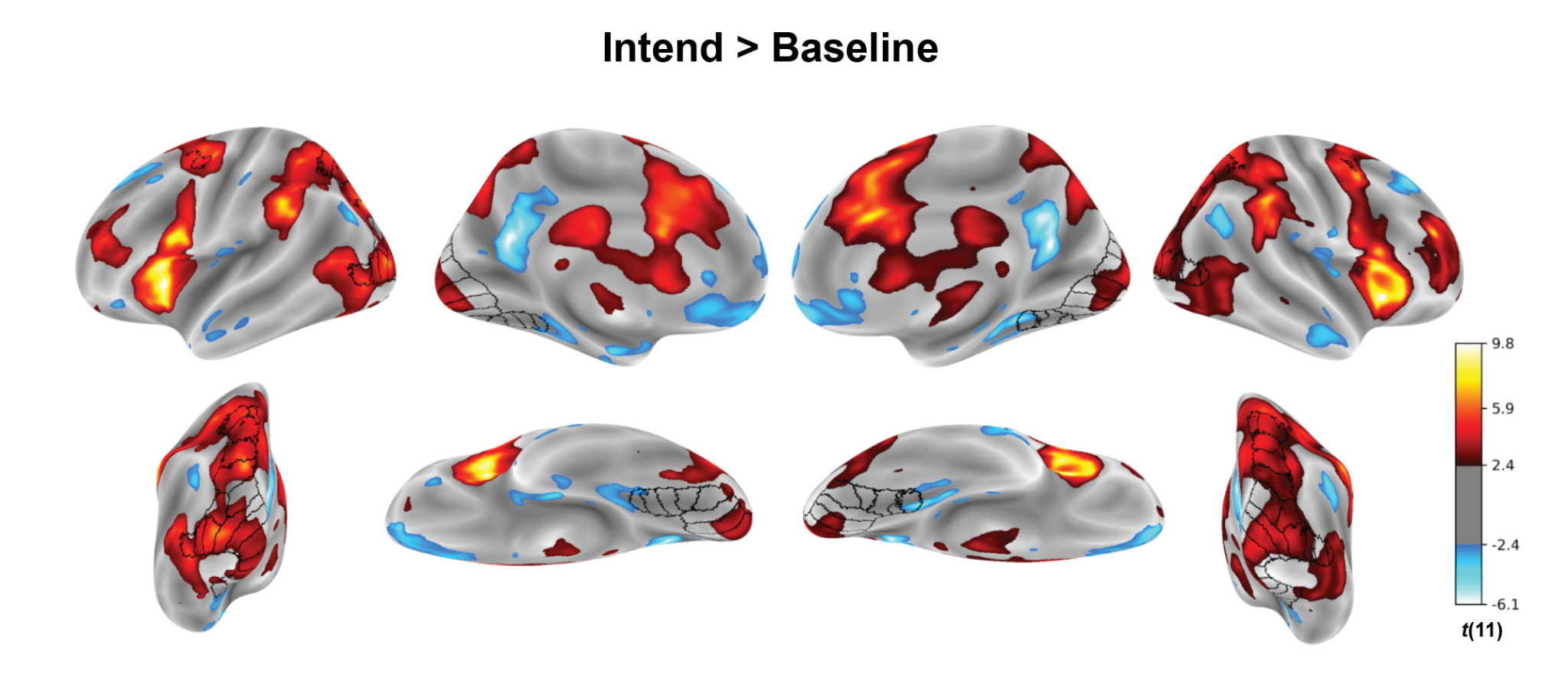

***Figure S2****.*

Group-level *t*-map showing the contrast between the intention and baseline periods in the volitional intention task (qFDR < 0.05). Black contours mark retinotopic visual regions defined using a probabilistic atlas (Wang et al., 2015).

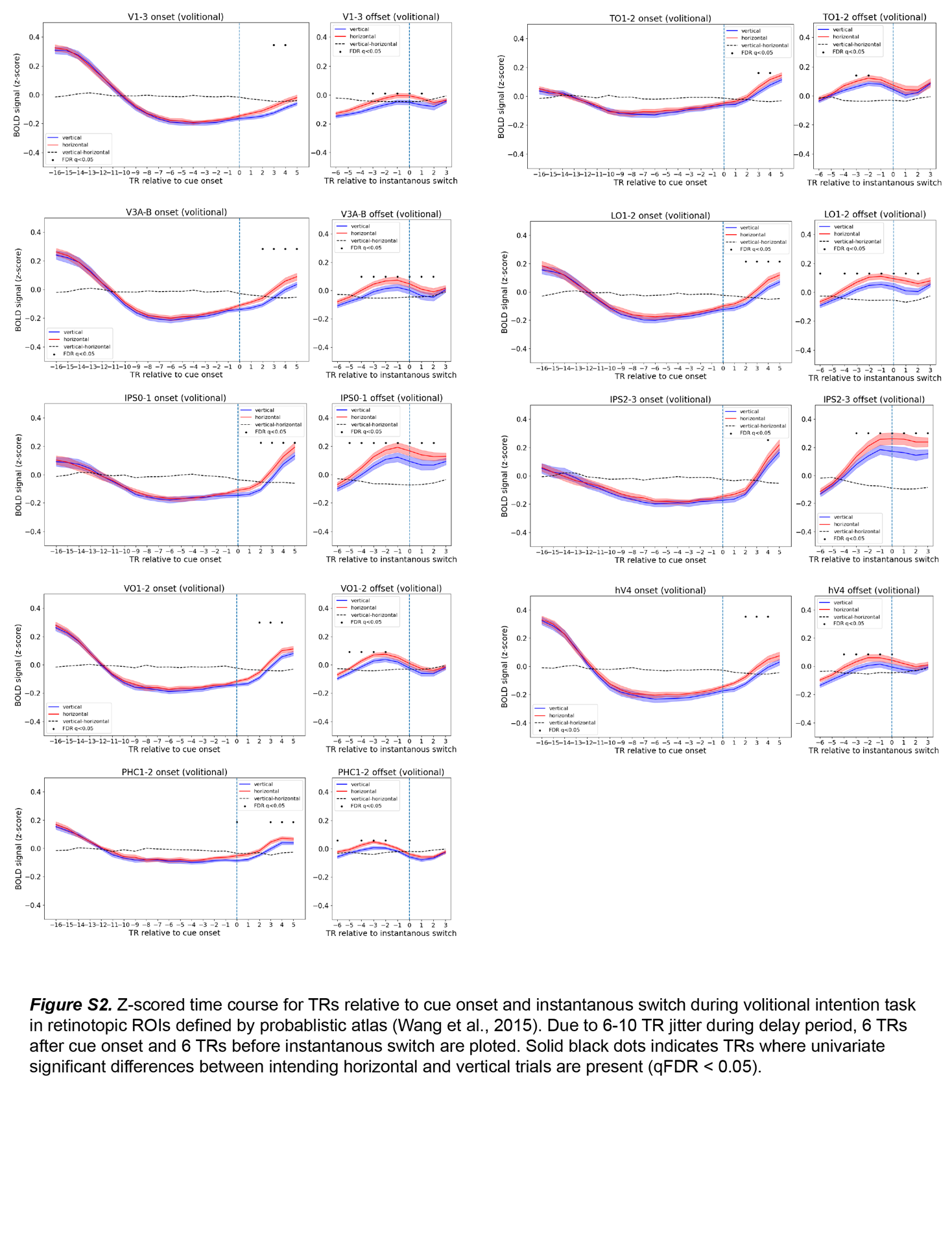

***Figure S3.***

Z-scored time course for TRs relative to cue onset and instantaneous switch during volitional intention task in retinotopic ROIs defined by probabilistic atlas (Wang et al., 2015). Due to 6-10 TR jitter during delay period, 6 TRs after cue onset and 6 TRs before instantaneous switch are plotted. Solid black dots indicates TRs where univariate significant differences between intending horizontal and vertical trials are present (qFDR < 0.05).

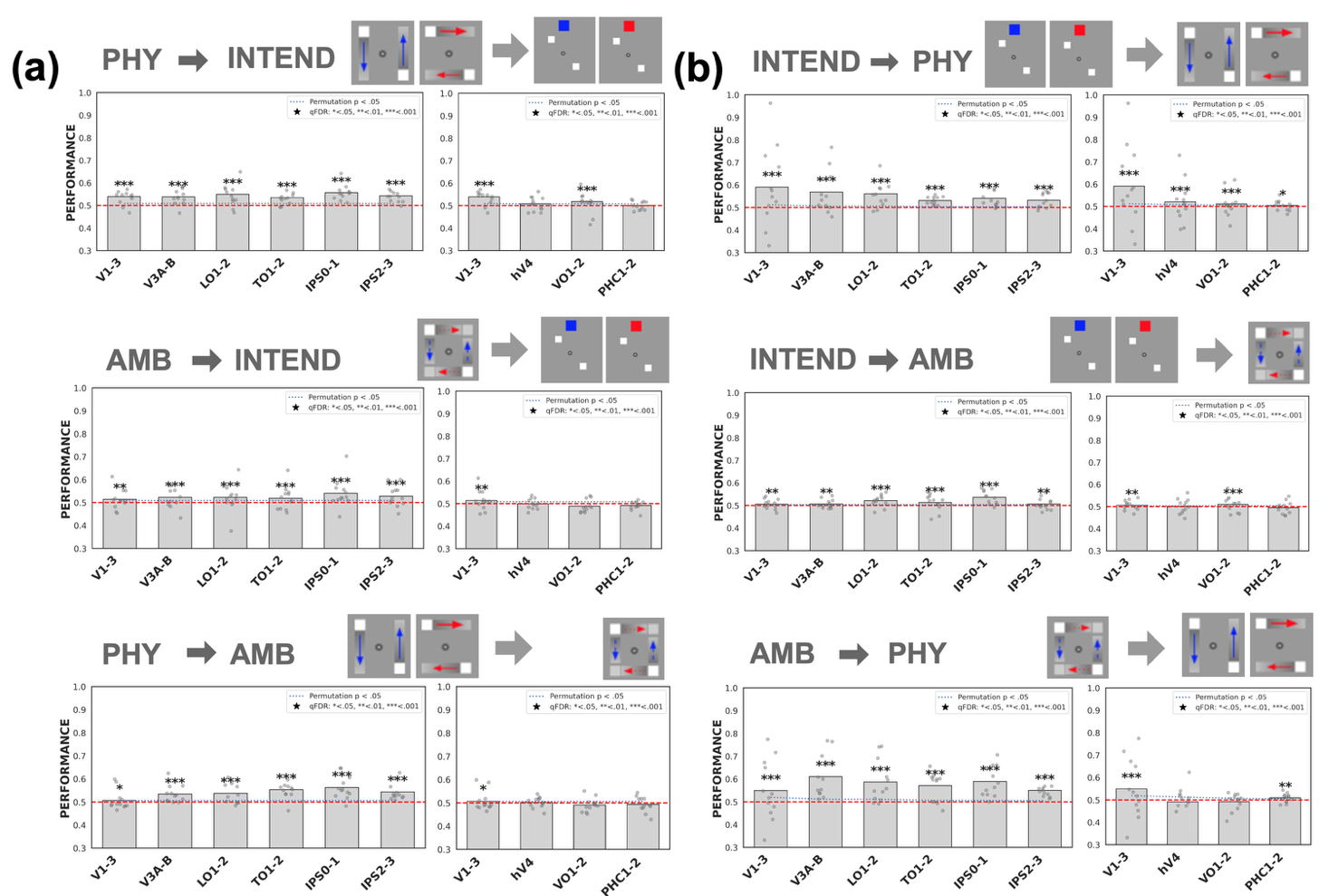

***Figure S4.***

Cross-decoding in Dorsal/Lateral and Ventral ROIs for both forward and backward train-test pair. SVM classifiers (a) forward cross-decoding (same as Figure 4a) and (b) backward cross-decoding.

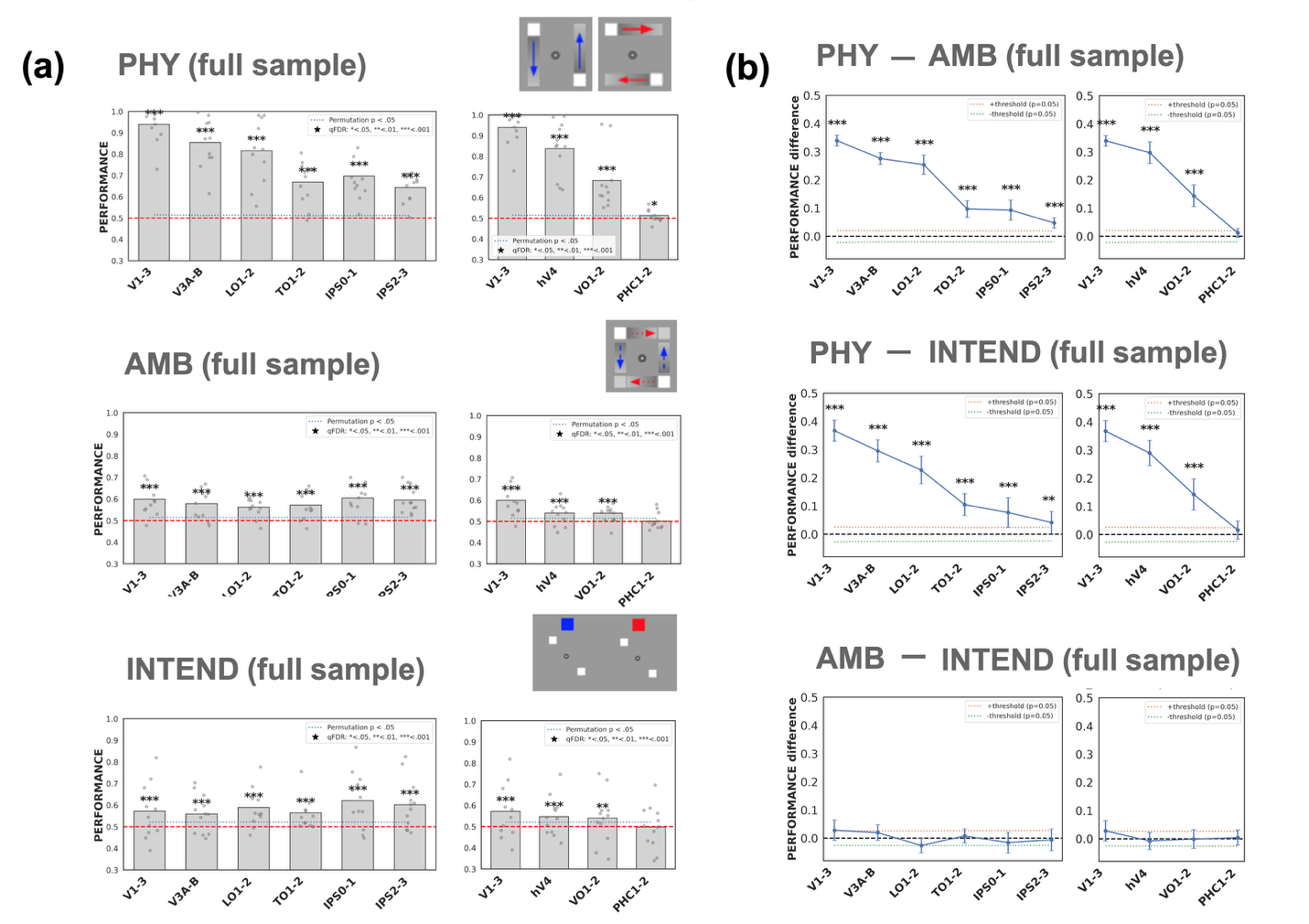

***Figure S5.***

Representational strength of veridical perception, illusion, and intention in dorsal and ventral ROIs using full samples***.*** (a) Leave-one-run-out decoding performance (AUC–ROC) for motion axis (horizontal vs. vertical) within physical (PHY), ambiguous (AMB), and volitional intention (INTEND) conditions. (b) Pairwise differences in decoding performance between conditions (PHY – AMB, PHY – INTEND, and AMB – INTEND). Plot conventions, permutation testing procedures, and FDR correction are identical to those described in Figure 5. Using full samples produced no qualitative differences compared to analyses using sub-sampled data (Figure 5).

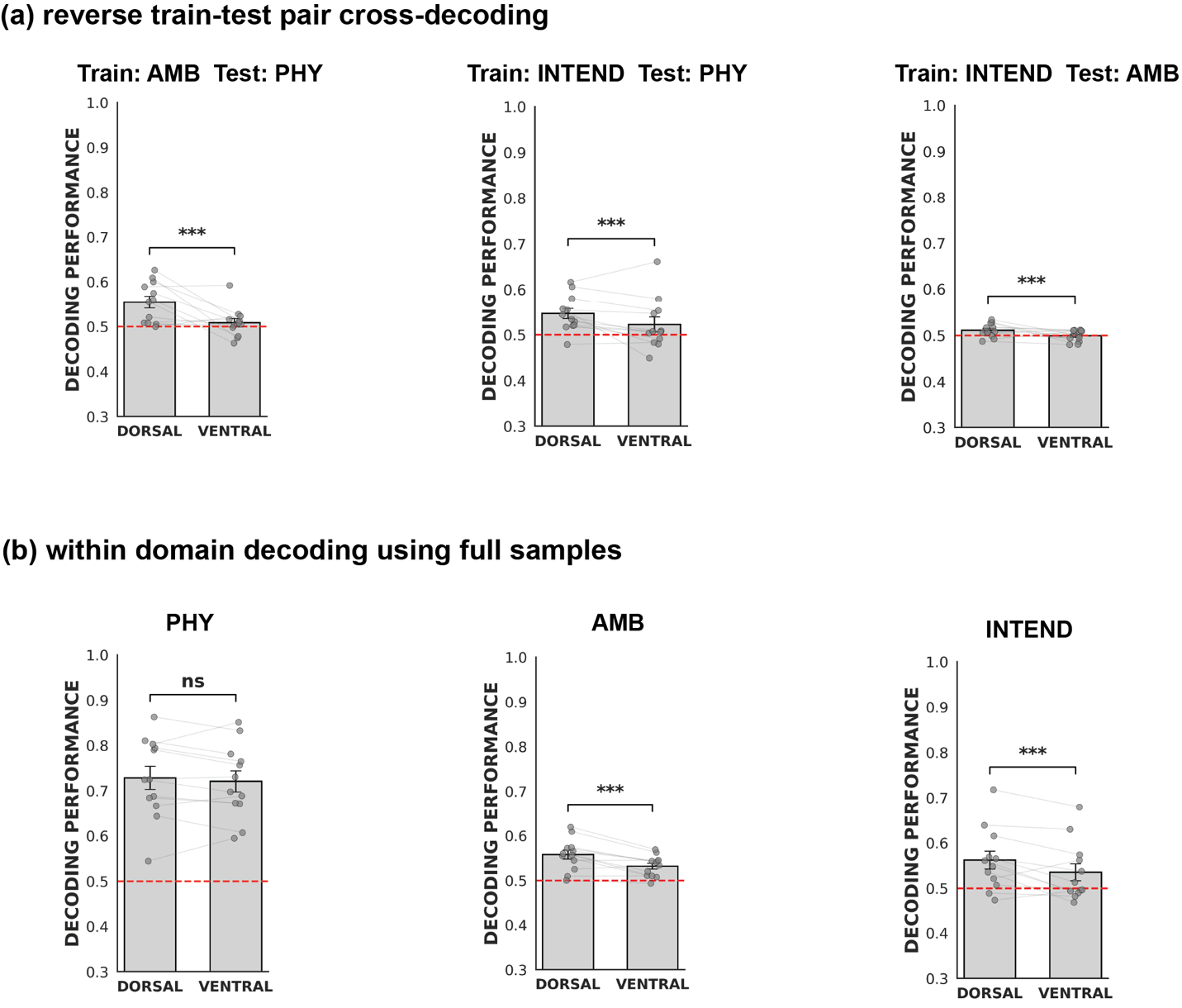

***Figure S6.***

a) Cross-condition decoding using the reverse train–test direction for each condition pair: classifiers were trained on ambiguous motion (AMB) and tested on physical motion (PHY), trained on volitional intention (INTEND) and tested on PHY, or trained on INTEND and tested on AMB. (b) Within-condition decoding of PHY, AMB, and INTEND using the full available samples. Bars show mean decoding performance across participants for dorsal/lateral and ventral visual ROIs; error bars indicate SEM. Gray points indicate individual participants, with lines connecting measurements from the same participant. The red dashed line indicates chance-level performance (0.5). Brackets indicate paired comparisons between dorsal/lateral and ventral decoding performance. Significance was assessed using paired permutation tests and corrected for multiple comparisons using the Benjamini–Hochberg false discovery rate (FDR) procedure. ****q* < 0.001; *ns*, not significant.

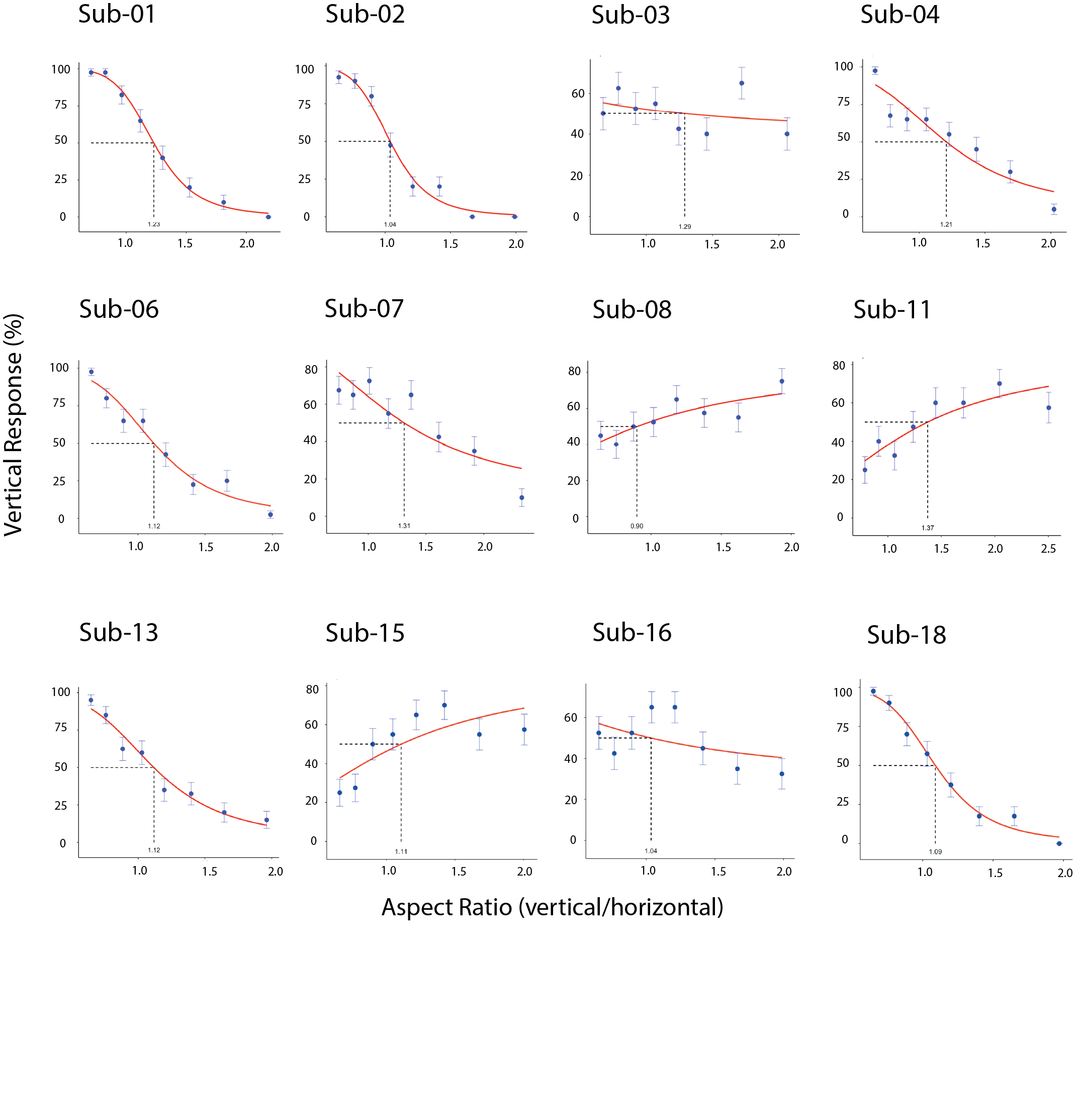

***Figure S7.* Subject-specific calibration of the motion quartet aspect ratio.**
Psychometric functions relating quartet aspect ratio to the proportion of vertical-motion percepts are shown for each participant. Before scanning, aspect ratio was varied across a fixed range to estimate a subject-specific point of subjective equality (PSE), defined as the aspect ratio associated with approximately equal reports of vertical and horizontal apparent motion (50% vertical responses; dashed lines). For several participants, perceptual reports were relatively insensitive to changes in aspect ratio over the range tested, resulting in shallow or poorly constrained psychometric fits. Importantly, the purpose of this procedure was not to characterize individual psychometric functions per se, but to calibrate the ambiguous stimulus used during scanning. The selected subject-specific aspect ratio provided a stimulus for which vertical and horizontal motion percepts occurred approximately equally often during passive viewing, thereby minimizing a systematic perceptual bias toward either motion axis. Points indicate observed response proportions [± error bars, standard errors of the mean], and red curves indicate fitted psychometric functions.

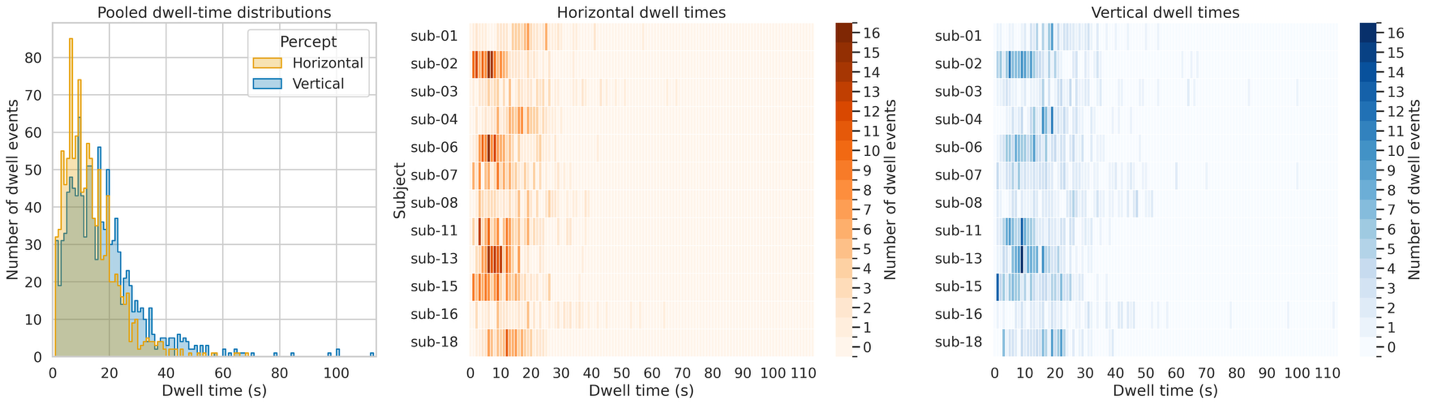

***Figure S8.***

Ambiguous apparent motion dwell time in the scanner

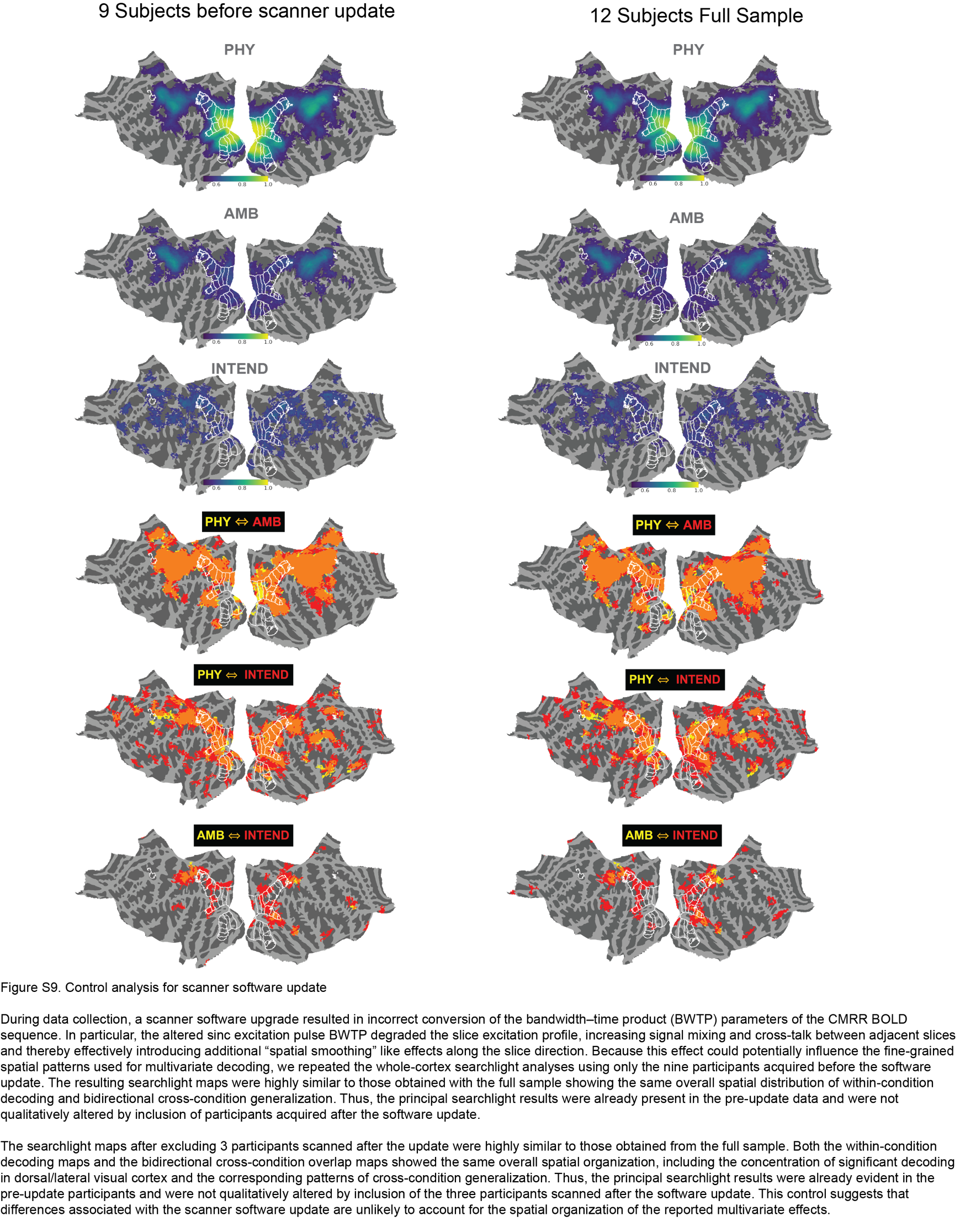

***Figure S9. Control analysis for scanner software update error***

During data collection, a scanner software upgrade resulted in incorrect conversion of the bandwidth–time product (BWTP) parameters of the BOLD sequence. In particular, the altered sinc excitation pulse BWTP degraded the slice excitation profile, increasing signal mixing and cross-talk between adjacent slices and thereby effectively introducing additional “spatial smoothing”-like effects along the slice direction. Because this effect could potentially influence the fine-grained spatial patterns used for multivariate decoding, we repeated the whole-cortex searchlight analyses using only the nine participants acquired before the software update. The resulting searchlight maps were highly similar to those obtained with the full sample showing the same overall spatial distribution of within-condition decoding and bidirectional cross-condition generalization. Thus, the principal searchlight results were already present in the pre-update data and were not qualitatively altered by inclusion of participants acquired after the software update.

The searchlight maps after excluding 3 participants scanned after the update were highly similar to those obtained from the full sample. Both the within-condition decoding maps and the bidirectional cross-condition overlap maps showed the same overall spatial organization, including the concentration of significant decoding in dorsal/lateral visual cortex and the corresponding patterns of cross-condition generalization. Thus, the principal searchlight results were already evident in the pre-update participants and were not qualitatively altered by inclusion of the three participants scanned after the software update. This control suggests that differences associated with the scanner software update are unlikely to account for the spatial organization of the reported multivariate effects.

**Table 1. Functionally defined hMT+ and atlas-defined hMT (TO1-2) decoding and cross-decoding performance**

| **Train** | **Test** | **ROI** | **ROC_AUC (MEAN ± SE)** | **Permutation p** | **FDR-adjusted p** | **N above FDR Threshold /12** | **Significant** |
| --- | --- | --- | --- | --- | --- | --- | --- |
| PHY | PHY | hMT+ | .72859 ± .03678 | < .00001 | .00001 | 11 | Yes |
| PHY | PHY | TO1-2 | .66927 ± .02861 | < .00001 | .00001 | 11 | Yes |
| PHY | AMB | hMT+ | .57311 ± .02028 | < .00001 | .00001 | 10 | Yes |
| PHY | AMB | TO1-2 | .55329 ± .01065 | < .00001 | .00001 | 11 | Yes |
| PHY | INTEND | hMT+ | .54179 ± .01199 | < .00001 | .00001 | 9 | Yes |
| PHY | INTEND | TO1-2 | .53477 ± .00828 | < .00001 | .00001 | 9 | Yes |
| AMB | PHY | hMT+ | .61352 ± .02531 | < .00001 | .00001 | 11 | Yes |
| AMB | PHY | TO1-2 | .57128 ± .01844 | < .00001 | .00001 | 8 | Yes |
| AMB | AMB | hMT+ | .59909 ± .02248 | < .00001 | .00001 | 9 | Yes |
| AMB | AMB | TO1-2 | .57227 ± .01861 | < .00001 | .00001 | 9 | Yes |
| AMB | INTEND | hMT+ | .52279 ± .01507 | < .00001 | .00001 | 7 | Yes |
| AMB | INTEND | TO1-2 | .51868 ± .01481 | .00048 | .00054 | 6 | Yes |
| INTEND | PHY | hMT+ | .55450 ± .01315 | < .00001 | .00001 | 10 | Yes |
| INTEND | PHY | TO1-2 | .53054 ± .00507 | < .00001 | .00001 | 12 | Yes |
| INTEND | AMB | hMT+ | .50736 ± .00929 | .00077 | .00081 | 7 | Yes |
| INTEND | AMB | TO1-2 | .51312 ± .01067 | < .00001 | .00001 | 8 | Yes |
| INTEND | INTEND | hMT+ | .52883 ± .01974 | .01253 | .01253 | 7 | Yes |
| INTEND | INTEND | TO1-2 | .56439 ± .02100 | < .00001 | .00001 | 7 | Yes |

**Table 2. Trial-balanced Wang atlas decoding and cross-decoding performance**

| **Train** | **Test** | **ROI** | **ROC_AUC (MEAN ± SE)** | **Permutation p** | **FDR-adjusted p** | **N above FDR Threshold /12** | **Significant** |
| --- | --- | --- | --- | --- | --- | --- | --- |
| AMB | AMB | V1-3 | .61963 ± .02079 | < .00001 | .00002 | 11 | Yes |
| AMB | AMB | V3A-B | .56644 ± .01964 | < .00001 | .00002 | 8 | Yes |
| AMB | AMB | LO1-2 | .55013 ± .01487 | < .00001 | .00002 | 8 | Yes |
| AMB | AMB | TO1-2 | .57923 ± .01595 | < .00001 | .00002 | 11 | Yes |
| AMB | AMB | IPS0-1 | .59107 ± .02664 | < .00001 | .00002 | 9 | Yes |
| AMB | AMB | IPS2-3 | .58906 ± .02338 | < .00001 | .00002 | 10 | Yes |
| AMB | AMB | hV4 | .53619 ± .01033 | .00100 | .00160 | 7 | Yes |
| AMB | AMB | VO1-2 | .55218 ± .01609 | .00003 | .00005 | 10 | Yes |
| AMB | AMB | PHC1-2 | .53329 ± .01548 | .00258 | .00397 | 7 | Yes |
| AMB | PHY | V1-3 | .54296 ± .02956 | .00052 | .00085 | 6 | Yes |
| AMB | PHY | V3A-B | .60328 ± .02516 | < .00001 | .00002 | 11 | Yes |
| AMB | PHY | LO1-2 | .56248 ± .02728 | < .00001 | .00002 | 8 | Yes |
| AMB | PHY | TO1-2 | .58531 ± .02035 | < .00001 | .00002 | 9 | Yes |
| AMB | PHY | IPS0-1 | .58241 ± .01798 | < .00001 | .00002 | 10 | Yes |
| AMB | PHY | IPS2-3 | .52359 ± .01174 | < .00001 | .00002 | 9 | Yes |
| AMB | PHY | hV4 | .48778 ± .02223 | .89852 | .92950 | 4 | No |
| AMB | PHY | VO1-2 | .49437 ± .01533 | .81225 | .86257 | 7 | No |
| AMB | PHY | PHC1-2 | .50734 ± .00793 | .05536 | .07549 | 6 | No |
| AMB | INTEND | V1-3 | .51177 ± .01380 | .02111 | .03016 | 5 | Yes |
| AMB | INTEND | V3A-B | .53699 ± .00690 | < .00001 | .00002 | 11 | Yes |
| AMB | INTEND | LO1-2 | .52597 ± .01533 | .00002 | .00004 | 8 | Yes |
| AMB | INTEND | TO1-2 | .51109 ± .01541 | .02426 | .03425 | 6 | Yes |
| AMB | INTEND | IPS0-1 | .54222 ± .01804 | < .00001 | .00002 | 8 | Yes |
| AMB | INTEND | IPS2-3 | .52475 ± .01328 | .00003 | .00005 | 8 | Yes |
| AMB | INTEND | hV4 | .49412 ± .00942 | .84714 | .89173 | 5 | No |
| AMB | INTEND | VO1-2 | .49917 ± .01200 | .55248 | .62545 | 4 | No |
| AMB | INTEND | PHC1-2 | .49772 ± .00857 | .67528 | .74343 | 3 | No |
| PHY | AMB | V1-3 | .50396 ± .01385 | .23192 | .28398 | 4 | No |
| PHY | AMB | V3A-B | .53342 ± .00991 | < .00001 | .00002 | 9 | Yes |
| PHY | AMB | LO1-2 | .54477 ± .01150 | < .00001 | .00002 | 9 | Yes |
| PHY | AMB | TO1-2 | .56037 ± .01159 | < .00001 | .00002 | 11 | Yes |
| PHY | AMB | IPS0-1 | .55523 ± .01659 | < .00001 | .00002 | 9 | Yes |
| PHY | AMB | IPS2-3 | .53625 ± .01108 | < .00001 | .00002 | 9 | Yes |
| PHY | AMB | hV4 | .49898 ± .00809 | .57561 | .64555 | 5 | No |
| PHY | AMB | VO1-2 | .49152 ± .00932 | .95483 | .97101 | 5 | No |
| PHY | AMB | PHC1-2 | .51676 ± .01120 | .00034 | .00058 | 6 | Yes |
| PHY | PHY | V1-3 | .94140 ± .02528 | < .00001 | .00002 | 12 | Yes |
| PHY | PHY | V3A-B | .84803 ± .03594 | < .00001 | .00002 | 12 | Yes |
| PHY | PHY | LO1-2 | .81269 ± .04380 | < .00001 | .00002 | 12 | Yes |
| PHY | PHY | TO1-2 | .65368 ± .03033 | < .00001 | .00002 | 12 | Yes |
| PHY | PHY | IPS0-1 | .65898 ± .03316 | < .00001 | .00002 | 10 | Yes |
| PHY | PHY | IPS2-3 | .62824 ± .02270 | < .00001 | .00002 | 9 | Yes |
| PHY | PHY | hV4 | .82851 ± .03759 | < .00001 | .00002 | 12 | Yes |
| PHY | PHY | VO1-2 | .66710 ± .03932 | < .00001 | .00002 | 12 | Yes |
| PHY | PHY | PHC1-2 | .48695 ± .01393 | .87291 | .91086 | 2 | No |
| PHY | INTEND | V1-3 | .52513 ± .00867 | < .00001 | .00002 | 7 | Yes |
| PHY | INTEND | V3A-B | .53141 ± .00909 | < .00001 | .00002 | 8 | Yes |
| PHY | INTEND | LO1-2 | .54302 ± .01615 | < .00001 | .00002 | 9 | Yes |
| PHY | INTEND | TO1-2 | .51436 ± .01000 | .00460 | .00690 | 8 | Yes |
| PHY | INTEND | IPS0-1 | .54893 ± .01248 | < .00001 | .00002 | 11 | Yes |
| PHY | INTEND | IPS2-3 | .54036 ± .01478 | < .00001 | .00002 | 9 | Yes |
| PHY | INTEND | hV4 | .49752 ± .00925 | .66934 | .74343 | 4 | No |
| PHY | INTEND | VO1-2 | .50631 ± .01023 | .11121 | .14994 | 7 | No |
| PHY | INTEND | PHC1-2 | .49026 ± .00796 | .96478 | .97289 | 2 | No |
| INTEND | AMB | V1-3 | .50325 ± .00903 | .14413 | .18206 | 5 | No |
| INTEND | AMB | V3A-B | .50371 ± .00702 | .12781 | .16491 | 5 | No |
| INTEND | AMB | LO1-2 | .51885 ± .00994 | < .00001 | .00002 | 8 | Yes |
| INTEND | AMB | TO1-2 | .50068 ± .01113 | .43563 | .50583 | 6 | No |
| INTEND | AMB | IPS0-1 | .53229 ± .00920 | < .00001 | .00002 | 8 | Yes |
| INTEND | AMB | IPS2-3 | .50219 ± .00661 | .24907 | .30190 | 4 | No |
| INTEND | AMB | hV4 | .49800 ± .01340 | .73576 | .79542 | 7 | No |
| INTEND | AMB | VO1-2 | .50986 ± .01192 | .00104 | .00164 | 7 | Yes |
| INTEND | AMB | PHC1-2 | .50085 ± .00749 | .37653 | .44297 | 6 | No |
| INTEND | PHY | V1-3 | .59874 ± .04695 | < .00001 | .00002 | 9 | Yes |
| INTEND | PHY | V3A-B | .56326 ± .02603 | < .00001 | .00002 | 9 | Yes |
| INTEND | PHY | LO1-2 | .56259 ± .01800 | < .00001 | .00002 | 11 | Yes |
| INTEND | PHY | TO1-2 | .53314 ± .00869 | < .00001 | .00002 | 10 | Yes |
| INTEND | PHY | IPS0-1 | .55083 ± .01138 | < .00001 | .00002 | 11 | Yes |
| INTEND | PHY | IPS2-3 | .53526 ± .01050 | < .00001 | .00002 | 10 | Yes |
| INTEND | PHY | hV4 | .51768 ± .02578 | .00031 | .00054 | 6 | Yes |
| INTEND | PHY | VO1-2 | .50685 ± .01595 | .03792 | .05230 | 4 | No |
| INTEND | PHY | PHC1-2 | .49259 ± .00927 | .99301 | .99301 | 5 | No |
| INTEND | INTEND | V1-3 | .57608 ± .03211 | < .00001 | .00002 | 8 | Yes |
| INTEND | INTEND | V3A-B | .56935 ± .02155 | < .00001 | .00002 | 7 | Yes |
| INTEND | INTEND | LO1-2 | .58042 ± .02947 | < .00001 | .00002 | 8 | Yes |
| INTEND | INTEND | TO1-2 | .55019 ± .02355 | .00010 | .00018 | 6 | Yes |
| INTEND | INTEND | IPS0-1 | .59945 ± .03784 | < .00001 | .00002 | 9 | Yes |
| INTEND | INTEND | IPS2-3 | .60124 ± .03465 | < .00001 | .00002 | 8 | Yes |
| INTEND | INTEND | hV4 | .51545 ± .02127 | .11869 | .15481 | 5 | No |
| INTEND | INTEND | VO1-2 | .53367 ± .03532 | .00514 | .00762 | 5 | Yes |
| INTEND | INTEND | PHC1-2 | .50645 ± .02694 | .30821 | .36985 | 5 | No |
| INTEND | AMB | V1-3 | .50370 ± .00924 | .11491 | .15153 | 5 | No |
| INTEND | AMB | V3A-B | .50026 ± .01031 | .47204 | .53948 | 5 | No |
| INTEND | AMB | LO1-2 | .52116 ± .01059 | < .00001 | .00002 | 8 | Yes |
| INTEND | AMB | TO1-2 | .50594 ± .01357 | .03083 | .04302 | 6 | Yes |
| INTEND | AMB | IPS0-1 | .53323 ± .01112 | < .00001 | .00002 | 8 | Yes |
| INTEND | AMB | IPS2-3 | .50044 ± .00700 | .43839 | .50583 | 4 | No |
| INTEND | AMB | hV4 | .49784 ± .01414 | .77504 | .83040 | 7 | No |
| INTEND | AMB | VO1-2 | .50861 ± .01383 | .00379 | .00576 | 7 | Yes |
| INTEND | AMB | PHC1-2 | .49830 ± .01015 | .71119 | .77585 | 6 | No |
| INTEND | PHY | V1-3 | .60650 ± .04789 | < .00001 | .00002 | 9 | Yes |
| INTEND | PHY | V3A-B | .56800 ± .02676 | < .00001 | .00002 | 9 | Yes |
| INTEND | PHY | LO1-2 | .56893 ± .01798 | < .00001 | .00002 | 11 | Yes |
| INTEND | PHY | TO1-2 | .53627 ± .00932 | < .00001 | .00002 | 10 | Yes |
| INTEND | PHY | IPS0-1 | .55180 ± .01252 | < .00001 | .00002 | 11 | Yes |
| INTEND | PHY | IPS2-3 | .54125 ± .01249 | < .00001 | .00002 | 10 | Yes |
| INTEND | PHY | hV4 | .51714 ± .02666 | .00058 | .00094 | 6 | Yes |
| INTEND | PHY | VO1-2 | .50297 ± .01699 | .22001 | .27501 | 4 | No |
| INTEND | PHY | PHC1-2 | .49533 ± .01181 | .94018 | .96429 | 5 | No |

*Note. N = 12 for all analyses. M = mean decoding ROC_AUC; SE = standard error. decimal-valued statistics are reported to five decimal places. FDR = false discovery rate. Significant indicates FDR q < .05.*

**Table 3. Full-sample decoding and cross-decoding performance (Figure 3)**

| **Train** | **Test** | **ROI** | **ROC_AUC (MEAN ± SE)** | **Permutation p** | **FDR-adjusted p** | **N above FDR threshold / 12** | **Significant** |
| --- | --- | --- | --- | --- | --- | --- | --- |
| AMB | AMB | V1-3 | .60000 ± .02000 | < .00001 | .00002 | 11 | Yes |
| AMB | AMB | V3A-B | .57900 ± .02000 | < .00001 | .00002 | 10 | Yes |
| AMB | AMB | LO1-2 | .56300 ± .01300 | < .00001 | .00002 | 10 | Yes |
| AMB | AMB | TO1-2 | .57200 ± .01900 | < .00001 | .00002 | 9 | Yes |
| AMB | AMB | IPS0-1 | .60500 ± .02000 | < .00001 | .00002 | 10 | Yes |
| AMB | AMB | IPS2-3 | .59600 ± .01700 | < .00001 | .00002 | 12 | Yes |
| AMB | AMB | hV4 | .53900 ± .01600 | < .00001 | .00002 | 8 | Yes |
| AMB | AMB | VO1-2 | .53900 ± .01200 | .00002 | .00003 | 8 | Yes |
| AMB | AMB | PHC1-2 | .50200 ± .01100 | .41800 | .47025 | 3 | No |
| AMB | PHY | V1-3 | .55000 ± .03700 | .00005 | .00007 | 6 | Yes |
| AMB | PHY | V3A-B | .61000 ± .02700 | < .00001 | .00002 | 11 | Yes |
| AMB | PHY | LO1-2 | .58700 ± .02700 | < .00001 | .00002 | 9 | Yes |
| AMB | PHY | TO1-2 | .57100 ± .01800 | < .00001 | .00002 | 8 | Yes |
| AMB | PHY | IPS0-1 | .59000 ± .01900 | < .00001 | .00002 | 11 | Yes |
| AMB | PHY | IPS2-3 | .55000 ± .00800 | < .00001 | .00002 | 12 | Yes |
| AMB | PHY | hV4 | .49200 ± .02200 | .80874 | .86651 | 4 | No |
| AMB | PHY | VO1-2 | .49300 ± .01100 | .93001 | .96959 | 5 | No |
| AMB | PHY | PHC1-2 | .50900 ± .00600 | .00152 | .00207 | 7 | Yes |
| AMB | INTEND | V1-3 | .51400 ± .01300 | .00498 | .00622 | 6 | Yes |
| AMB | INTEND | V3A-B | .52300 ± .01300 | .00003 | .00005 | 6 | Yes |
| AMB | INTEND | LO1-2 | .52300 ± .01800 | .00005 | .00007 | 8 | Yes |
| AMB | INTEND | TO1-2 | .51900 ± .01500 | .00048 | .00068 | 6 | Yes |
| AMB | INTEND | IPS0-1 | .54000 ± .01800 | < .00001 | .00002 | 9 | Yes |
| AMB | INTEND | IPS2-3 | .52800 ± .01300 | < .00001 | .00002 | 7 | Yes |
| AMB | INTEND | hV4 | .50000 ± .00700 | .50633 | .55573 | 4 | No |
| AMB | INTEND | VO1-2 | .48900 ± .00800 | .98185 | .98521 | 3 | No |
| AMB | INTEND | PHC1-2 | .49200 ± .00600 | .93335 | .96959 | 1 | No |
| PHY | AMB | V1-3 | .50700 ± .01200 | .03648 | .04320 | 4 | Yes |
| PHY | AMB | V3A-B | .53500 ± .01100 | < .00001 | .00002 | 9 | Yes |
| PHY | AMB | LO1-2 | .53800 ± .01300 | < .00001 | .00002 | 7 | Yes |
| PHY | AMB | TO1-2 | .55300 ± .01100 | < .00001 | .00002 | 11 | Yes |
| PHY | AMB | IPS0-1 | .56300 ± .01400 | < .00001 | .00002 | 11 | Yes |
| PHY | AMB | IPS2-3 | .54300 ± .01100 | < .00001 | .00002 | 12 | Yes |
| PHY | AMB | hV4 | .50300 ± .00700 | .24889 | .28718 | 6 | No |
| PHY | AMB | VO1-2 | .49100 ± .00800 | .98521 | .98521 | 2 | No |
| PHY | AMB | PHC1-2 | .49400 ± .01000 | .93727 | .96959 | 5 | No |
| PHY | PHY | V1-3 | .94000 ± .02300 | < .00001 | .00002 | 12 | Yes |
| PHY | PHY | V3A-B | .85500 ± .03300 | < .00001 | .00002 | 12 | Yes |
| PHY | PHY | LO1-2 | .81600 ± .04100 | < .00001 | .00002 | 12 | Yes |
| PHY | PHY | TO1-2 | .66900 ± .02900 | < .00001 | .00002 | 11 | Yes |
| PHY | PHY | IPS0-1 | .69700 ± .03200 | < .00001 | .00002 | 12 | Yes |
| PHY | PHY | IPS2-3 | .64400 ± .01900 | < .00001 | .00002 | 11 | Yes |
| PHY | PHY | hV4 | .83700 ± .03700 | < .00001 | .00002 | 12 | Yes |
| PHY | PHY | VO1-2 | .68300 ± .03900 | < .00001 | .00002 | 12 | Yes |
| PHY | PHY | PHC1-2 | .51400 ± .00900 | .03324 | .04043 | 4 | Yes |
| PHY | INTEND | V1-3 | .54000 ± .01000 | < .00001 | .00002 | 10 | Yes |
| PHY | INTEND | V3A-B | .53900 ± .01000 | < .00001 | .00002 | 9 | Yes |
| PHY | INTEND | LO1-2 | .55000 ± .01500 | < .00001 | .00002 | 10 | Yes |
| PHY | INTEND | TO1-2 | .53500 ± .00800 | < .00001 | .00002 | 9 | Yes |
| PHY | INTEND | IPS0-1 | .55700 ± .01000 | < .00001 | .00002 | 12 | Yes |
| PHY | INTEND | IPS2-3 | .54400 ± .00800 | < .00001 | .00002 | 11 | Yes |
| PHY | INTEND | hV4 | .50800 ± .00900 | .05885 | .06878 | 6 | No |
| PHY | INTEND | VO1-2 | .51800 ± .01400 | .00027 | .00039 | 10 | Yes |
| PHY | INTEND | PHC1-2 | .50100 ± .00600 | .43457 | .48285 | 5 | No |
| INTEND | AMB | V1-3 | .50600 ± .00600 | .00469 | .00603 | 6 | Yes |
| INTEND | AMB | V3A-B | .50700 ± .00600 | .00278 | .00373 | 5 | Yes |
| INTEND | AMB | LO1-2 | .52100 ± .00700 | < .00001 | .00002 | 10 | Yes |
| INTEND | AMB | TO1-2 | .51300 ± .01100 | < .00001 | .00002 | 9 | Yes |
| INTEND | AMB | IPS0-1 | .53600 ± .00900 | < .00001 | .00002 | 9 | Yes |
| INTEND | AMB | IPS2-3 | .50700 ± .00800 | .00291 | .00385 | 5 | Yes |
| INTEND | AMB | hV4 | .50200 ± .01000 | .25926 | .29536 | 6 | No |
| INTEND | AMB | VO1-2 | .51000 ± .01000 | .00003 | .00005 | 7 | Yes |
| INTEND | AMB | PHC1-2 | .49500 ± .00800 | .98307 | .98521 | 5 | No |
| INTEND | PHY | V1-3 | .59100 ± .05000 | < .00001 | .00002 | 8 | Yes |
| INTEND | PHY | V3A-B | .56900 ± .02700 | < .00001 | .00002 | 9 | Yes |
| INTEND | PHY | LO1-2 | .56200 ± .01700 | < .00001 | .00002 | 10 | Yes |
| INTEND | PHY | TO1-2 | .53100 ± .00500 | < .00001 | .00002 | 12 | Yes |
| INTEND | PHY | IPS0-1 | .54000 ± .01000 | < .00001 | .00002 | 10 | Yes |
| INTEND | PHY | IPS2-3 | .53200 ± .00900 | < .00001 | .00002 | 9 | Yes |
| INTEND | PHY | hV4 | .52000 ± .02700 | .00004 | .00006 | 6 | Yes |
| INTEND | PHY | VO1-2 | .51100 ± .01600 | .00014 | .00020 | 7 | Yes |
| INTEND | PHY | PHC1-2 | .50400 ± .00700 | .03114 | .03839 | 7 | Yes |
| INTEND | INTEND | V1-3 | .57200 ± .03600 | < .00001 | .00002 | 7 | Yes |
| INTEND | INTEND | V3A-B | .55900 ± .02500 | .00003 | .00005 | 8 | Yes |
| INTEND | INTEND | LO1-2 | .58900 ± .02500 | < .00001 | .00002 | 10 | Yes |
| INTEND | INTEND | TO1-2 | .56400 ± .02100 | < .00001 | .00002 | 7 | Yes |
| INTEND | INTEND | IPS0-1 | .62000 ± .03700 | < .00001 | .00002 | 9 | Yes |
| INTEND | INTEND | IPS2-3 | .60200 ± .03300 | < .00001 | .00002 | 8 | Yes |
| INTEND | INTEND | hV4 | .54800 ± .02600 | .00006 | .00009 | 7 | Yes |
| INTEND | INTEND | VO1-2 | .54000 ± .03400 | .00104 | .00144 | 7 | Yes |
| INTEND | INTEND | PHC1-2 | .49800 ± .03000 | .55446 | .60123 | 5 | No |

Note. N = 12 for all analyses. ROC_AUC values are group mean ± standard error. Permutation p values and FDR-adjusted p values are reported from the supplied full-sample Figure 3 dataset.

**Table 4. Representational strength comparisons, trial-balanced analysis**

| **Condition** | **ROI** | **Mean difference ± SE** | **Permutation p** | **Negative threshold (p < .05)** | **Positive threshold (p < .05)** | **FDR-adjusted p** | **Significant** |
| --- | --- | --- | --- | --- | --- | --- | --- |
| AMB - INTEND | V1-3 | .04354 ± .02864 | .01364 | -.02941 | .02841 | .02352 | Yes |
| AMB - INTEND | V3A-B | -.00291 ± .02732 | .86939 | -.02889 | .02838 | .88713 | No |
| AMB - INTEND | LO1-2 | -.03029 ± .02403 | .08598 | -.02860 | .02950 | .13027 | No |
| AMB - INTEND | TO1-2 | .02904 ± .02525 | .09757 | -.02866 | .02902 | .14348 | No |
| AMB - INTEND | IPS0-1 | -.00838 ± .04504 | .63437 | -.02895 | .02888 | .66081 | No |
| AMB - INTEND | IPS2-3 | -.01218 ± .04499 | .49236 | -.02906 | .02901 | .53518 | No |
| PHY – AMB | V1-3 | .32177 ± .01988 | .00001 | -.02830 | .02938 | .00002 | Yes |
| PHY – AMB | V3A-B | .28159 ± .03129 | .00001 | -.02725 | .02683 | .00002 | Yes |
| PHY – AMB | LO1-2 | .26256 ± .04062 | .00001 | -.02718 | .02666 | .00002 | Yes |
| PHY – AMB | TO1-2 | .07445 ± .02311 | .00001 | -.02574 | .02546 | .00002 | Yes |
| PHY – AMB | IPS0-1 | .06791 ± .02867 | .00003 | -.02582 | .02689 | .00007 | Yes |
| PHY – AMB | IPS2-3 | .03918 ± .02648 | .01343 | -.02673 | .02536 | .02352 | Yes |
| PHY - INTEND | V1-3 | .36532 ± .03749 | .00001 | -.03166 | .03178 | .00002 | Yes |
| PHY - INTEND | V3A-B | .27868 ± .03593 | .00001 | -.03007 | .02887 | .00002 | Yes |
| PHY - INTEND | LO1-2 | .23227 ± .05220 | .00001 | -.02914 | .02912 | .00002 | Yes |
| PHY - INTEND | TO1-2 | .10349 ± .04443 | .00001 | -.02822 | .02809 | .00002 | Yes |
| PHY - INTEND | IPS0-1 | .05953 ± .05537 | .00057 | -.02802 | .02901 | .00124 | Yes |
| PHY - INTEND | IPS2-3 | .02700 ± .04010 | .11330 | -.02880 | .02722 | .16186 | No |
| AMB - INTEND | hV4 | .02074 ± .01776 | .23352 | -.02845 | .02905 | .29190 | No |
| AMB - INTEND | VO1-2 | .01851 ± .03812 | .29260 | -.03007 | .02773 | .34074 | No |
| AMB - INTEND | PHC1-2 | .02684 ± .02950 | .12887 | -.02935 | .02868 | .17415 | No |
| PHY - AMB | hV4 | .29232 ± .03585 | .00001 | -.02846 | .02730 | .00002 | Yes |
| PHY - AMB | VO1-2 | .11492 ± .03796 | .00001 | -.02601 | .02780 | .00002 | Yes |
| PHY - AMB | PHC1-2 | -.04634 ± .01870 | .00580 | -.02690 | .02834 | .01208 | Yes |
| PHY - INTEND | hV4 | .31306 ± .04273 | .00001 | -.03018 | .02983 | .00002 | Yes |
| PHY - INTEND | VO1-2 | .13343 ± .05320 | .00001 | -.02870 | .02792 | .00002 | Yes |
| PHY - INTEND | PHC1-2 | -.01949 ± .03339 | .26621 | -.02842 | .02918 | .32464 | No |

Note. N = 12 for all analyses. All decimal-valued statistics are reported to five decimal places. Negative and positive columns give the two-sided permutation thresholds at p < .05. Significant indicates FDR q < .05.

**Table 5. Representational strength comparisons, full sample**

| **Condition** | **ROI** | **Mean difference ± SE** | **Permutation p** | **Negative threshold (p < .05)** | **Positive threshold (p < .05)** | **FDR-adjusted p** | **Significant** |
| --- | --- | --- | --- | --- | --- | --- | --- |
| AMB – INTEND | V1-3 | .02780 ± .03631 | .08219 | -.02558 | .02687 | .12328 | No |
| AMB – INTEND | V3A-B | .01954 ± .02783 | .21613 | -.02616 | .02577 | .29472 | No |
| AMB – INTEND | LO1-2 | -.02613 ± .02632 | .09729 | -.02650 | .02533 | .13898 | No |
| AMB – INTEND | TO1-2 | .00788 ± .02524 | .61885 | -.02598 | .02604 | .68761 | No |
| AMB – INTEND | IPS0-1 | -.01588 ± .03685 | .32121 | -.02630 | .02646 | .38902 | No |
| AMB – INTEND | IPS2-3 | -.00607 ± .03880 | .70140 | -.02530 | .02681 | .75150 | No |
| PHY – AMB | V1-3 | .33950 ± .01913 | .00001 | -.02262 | .02038 | .00002 | Yes |
| PHY – AMB | V3A-B | .27612 ± .02151 | .00001 | -.02031 | .01979 | .00002 | Yes |
| PHY – AMB | LO1-2 | .25370 ± .03454 | .00001 | -.01977 | .02042 | .00002 | Yes |
| PHY – AMB | TO1-2 | .09700 ± .02947 | .00001 | -.02014 | .01886 | .00002 | Yes |
| PHY – AMB | IPS0-1 | .09274 ± .03552 | .00001 | -.02051 | .01934 | .00002 | Yes |
| PHY – AMB | IPS2-3 | .04764 ± .01842 | .00010 | -.01994 | .01887 | .00018 | Yes |
| PHY – INTEND | V1-3 | .36730 ± .03730 | .00001 | -.02736 | .02619 | .00002 | Yes |
| PHY – INTEND | V3A-B | .29566 ± .03929 | .00001 | -.02572 | .02486 | .00002 | Yes |
| PHY – INTEND | LO1-2 | .22757 ± .04894 | .00001 | -.02551 | .02515 | .00002 | Yes |
| PHY – INTEND | TO1-2 | .10488 ± .03865 | .00001 | -.02507 | .02381 | .00002 | Yes |
| PHY – INTEND | IPS0-1 | .07685 ± .05215 | .00001 | -.02489 | .02396 | .00002 | Yes |
| PHY – INTEND | IPS2-3 | .04157 ± .03949 | .00492 | -.02409 | .02481 | .00820 | Yes |
| AMB – INTEND | hV4 | -.00810 ± .02996 | .60808 | -.02566 | .02620 | .68761 | No |
| AMB – INTEND | VO1-2 | -.00128 ± .03331 | .93476 | -.02612 | .02617 | .93476 | No |
| AMB – INTEND | PHC1-2 | .00384 ± .02670 | .81027 | -.02620 | .02656 | .83821 | No |
| PHY – AMB | hV4 | .29734 ± .03888 | .00001 | -.02128 | .02015 | .00002 | Yes |
| PHY – AMB | VO1-2 | .14354 ± .03904 | .00001 | -.02091 | .01962 | .00002 | Yes |
| PHY – AMB | PHC1-2 | .01190 ± .01580 | .32419 | -.02018 | .01943 | .38902 | No |
| PHY – INTEND | hV4 | .28924 ± .04490 | .00001 | -.02584 | .02550 | .00002 | Yes |
| PHY – INTEND | VO1-2 | .14227 ± .05544 | .00001 | -.02577 | .02450 | .00002 | Yes |
| PHY – INTEND | PHC1-2 | .01574 ± .03235 | .29667 | -.02506 | .02459 | .38696 | No |

Note. N = 12 for all analyses. All decimal-valued statistics are reported to five decimal places. Negative and positive columns give the two-sided permutation thresholds at p < .05. Significant indicates FDR q < .05.

**Table 6 Dorsal – Ventral comparisons statistics (Full sample)**

| **trained on** | **target** | **Dorsal Mean** | **Dorsal SEM** | **Ventral Mean** | **Ventral SEM** | **Dorsal minus Ventral** | **Negative Permutation Threshold p05** | **Positive Permutation Threshold p05** | **Permutation p** | **FDR Adjusted p** | **FDR Significant q05** |
| --- | --- | --- | --- | --- | --- | --- | --- | --- | --- | --- | --- |
| PHY | PHY | 0.717407 | 0.024469 | 0.711644 | 0.023414 | 0.005763 | -9.0e-03 | 0.008818 | 0.289767 | 0.289767 | No |
| AMB | AMB | 0.559641 | 0.011673 | 0.541391 | 0.007882 | 0.018249 | -9.0e-03 | 0.008821 | 7.9e-04 | 8.6e-04 | Yes |
| INTEND | INTEND | 0.559063 | 0.019783 | 0.526263 | 0.017085 | 0.0328 | -1.0e-02 | 0.009998 | 1.0e-05 | 1.3e-05 | Yes |
| PHY | AMB | 0.530052 | 0.005461 | 0.500657 | 0.005357 | 0.029396 | -3.7e-03 | 0.003704 | 1.0e-05 | 1.3e-05 | Yes |
| AMB | PHY | 0.548445 | 0.012265 | 0.514188 | 0.009151 | 0.034256 | -7.6e-03 | 0.007388 | 1.0e-05 | 1.3e-05 | Yes |
| PHY | INTEND | 0.527311 | 0.003858 | 0.497119 | 0.005251 | 0.030192 | -4.1e-03 | 0.004318 | 1.0e-05 | 1.3e-05 | Yes |
| INTEND | PHY | 0.547674 | 0.011089 | 0.518711 | 0.015585 | 0.028963 | -4.2e-03 | 0.004273 | 1.0e-05 | 1.3e-05 | Yes |
| AMB | INTEND | 0.515321 | 0.007224 | 0.494353 | 0.003434 | 0.020968 | -4.3e-03 | 0.004376 | 1.0e-05 | 1.3e-05 | Yes |
| INTEND | AMB | 0.508771 | 0.004906 | 0.497107 | 0.005699 | 0.011664 | -2.4e-03 | 0.002358 | 1.0e-05 | 1.3e-05 | Yes |

**Table 7 Dorsal – Ventral comparisons statistics (Full sample)**

| **trained on** | **target** | **Dorsal Mean** | **Dorsal SEM** | **Ventral Mean** | **Ventral SEM** | **Dorsal minus Ventral** | **Negative Permutation Threshold p05** | **Positive Permutation Threshold p05** | **Permutation p** | **FDR Adjusted p** | **FDR Significant q05** |
| --- | --- | --- | --- | --- | --- | --- | --- | --- | --- | --- | --- |
| PHY | PHY | 0.727965 | 0.02563 | 0.720631 | 0.023259 | 0.007334 | -6.1e-03 | 0.00641 | 0.055219 | 0.055219 | No |
| AMB | AMB | 0.55785 | 0.010259 | 0.531827 | 0.00673 | 0.026023 | -7.1e-03 | 0.006667 | 1.0e-05 | 1.3e-05 | Yes |
| INTEND | INTEND | 0.562078 | 0.019765 | 0.535488 | 0.018749 | 0.026591 | -9.5e-03 | 0.009858 | 3.0e-05 | 3.4e-05 | Yes |
| PHY | AMB | 0.533846 | 0.005924 | 0.501564 | 0.003027 | 0.032281 | -2.9e-03 | 0.002955 | 1.0e-05 | 1.3e-05 | Yes |
| AMB | PHY | 0.554639 | 0.013104 | 0.508614 | 0.009513 | 0.046025 | -6.2e-03 | 0.006994 | 1.0e-05 | 1.3e-05 | Yes |
| PHY | INTEND | 0.534162 | 0.004227 | 0.506718 | 0.004669 | 0.027444 | -4.2e-03 | 0.003933 | 1.0e-05 | 1.3e-05 | Yes |
| vol | PHY | 0.546482 | 0.011359 | 0.522466 | 0.016229 | 0.024016 | -3.9e-03 | 0.00398 | 1.0e-05 | 1.3e-05 | Yes |
| AMB | INTEND | 0.515947 | 0.0077 | 0.496891 | 0.003635 | 0.019055 | -4.1e-03 | 0.004263 | 1.0e-05 | 1.3e-05 | Yes |
| vol | AMB | 0.511156 | 0.004195 | 0.499554 | 0.003456 | 0.011603 | -1.8e-03 | 0.001864 | 1.0e-05 | 1.3e-05 | Yes |
